# Subcortical Auditory Model including Efferent Dynamic Gain Control with Inputs from Cochlear Nucleus and Inferior Colliculus

**DOI:** 10.1101/2022.10.25.513794

**Authors:** Afagh Farhadi, Skyler G. Jennings, Elizabeth A. Strickland, Laurel H. Carney

## Abstract

We developed an auditory model with a time-varying, gain-control signal based on the physiology of the efferent system and the sub-cortical neural pathways. The medial olivocochlear (MOC) efferent stage of the model receives excitatory projections from both fluctuation-sensitive model neurons of the inferior colliculus (IC) and wide-dynamic-range model neurons of the cochlear nucleus. The response of the model MOC stage dynamically controls cochlear gain via simulated outer hair cells. In response to amplitude-modulated (AM) noise, firing rates of most IC neurons with band-enhanced modulation transfer functions in awake rabbits increase over a time course consistent with the dynamics of the MOC efferent feedback. These changes in the rates of IC neurons in awake rabbits were employed to adjust the parameters of the efferent stage of the proposed model. Responses of the proposed model to AM noise were able to simulate the increasing IC rate over time, while the model without the efferent system did not show this trend. The proposed model with efferent gain control provides a powerful tool for testing hypotheses, shedding insight on mechanisms in hearing, specifically those involving the efferent system.

## I. INTRODUCTION

The descending auditory pathway, or efferent system, is as large as the ascending pathway in terms of number of projections, but this part of the system not been as extensively studied as the ascending pathway. The efferent system consists of several feedback projections (Guinan, 2006; Warr, 1992) spanning all levels of the auditory pathway, from the auditory cortex to the cochlea. The focus of this study is the medial olivocochlear (MOC) system, which includes projections from MOC neurons to the outer hair cells and directly controls cochlear gain. Although several studies suggest the critical role of the MOC efferent system for interpreting data from psychoacoustic measures (Lopez-Poveda, 2018; Jennings, 2021; Lauer et al., 2022), it is still unclear the extent to which gain control by the MOC efferent system is involved in hearing. Understanding the role of the efferent system in auditory processing is essential to gain a comprehensive understanding of auditory function. In this study, we develop a computational model for the sub-cortical auditory system, providing a tool for exploring efferent mechanisms of cochlear gain control.

Despite several studies of the physiology of the efferent system (Guinan, 2006; Gummer et al., 1988; Liberman & Guinan, 1998), many aspects of this system are not well understood, such as the effects of anesthesia, the role of higher-level projections, and the independent effects of the MOC and middle-ear muscle reflexes. This gap in understanding is potentially explained by its complexity and the limited experimental methods for understanding the efferent system (Schofield, 2011). Most of the information available is based on a relatively small number of recordings from single neurons located deep in the brainstem in anesthetized animals (Guinan, 2018); however, anesthesia suppresses MOC responses (Aedo et al., 2015; Chambers et al., 2012; Guitton et al., 2004). Noninvasive methods used to study MOC effects on cochlear gain include measurements of either otoacoustic emissions (OAEs) (Guinan, 2006),the cochlear microphonic (Jennings & Aviles, 2023; Jamos et al., 2020, 2021), or the compound action potential (Lichtenhan et al. 2016; Smith et al., 2017), with and without MOC activation by contralateral broadband noise. These methods provide useful insights about the efferent system; however, they do not provide details about the contributions of higher-level projections in the pathway, such as those from the IC or auditory cortex to the MOC (Delano et al, 2007). Furthermore, care must be taken to separate effects of the middle-ear muscle reflex on OAEs when studying the MOC efferent system (Sun, 2008, Guinan, 2018). Here we propose a model of the MOC efferent system that can help in designing future physiological and psychophysical experiments for studying efferent function.

A new framework for auditory coding proposes an explicit role for the efferent system. Carney (2018) suggests that low-frequency temporal fluctuations in auditory-nerve (AN) responses can be a reliable cue for auditory perception. It is often assumed that AN average-rate profiles encode the spectrum of complex sounds; however, the majority of AN fibers, the high-spontaneous-rate (HSR) fibers, have average rates that are saturated at conversational speech levels (55–65 dB SPL, Olsen, 1998). Therefore, in many conditions, average discharge rates of HSR AN fibers do not contain enough information to explain perception of complex sounds (Bharadwaj et al., 2014; Carney, 2018; Liberman, 1978). On the other hand, temporal fluctuations of HSR AN responses, referred to as neural fluctuations, have robust information at conversational speech levels, potentially encoding peaks and valleys in the stimulus spectrum because the fluctuation depths are smaller for AN fibers tuned near peak frequencies and larger for other frequency channels (Carney et al., 2015; Carney, 2018). If neural fluctuations play a key role in encoding complex sounds, then these fluctuations may be involved in the efferent gain-control system to maintain contrasts in the neural-fluctuation pro-file. Interestingly, neurons in the auditory midbrain (inferior colliculus, IC) are sensitive to low-frequency (i.e., < 100Hz) fluctuations of their inputs (Joris et al. 2004; Krishna & Semple, 2000), and the IC projects to MOC neurons (Huffman & Henson 1990; Delano et al., 2007).

The MOC efferent system receives several convergent inputs; the model presented here focuses on subcortical pathways, specifically inputs from the IC and cochlear nucleus (CN). MOC neurons receive descending projections from IC cells in the midbrain and exhibit band-pass modulation transfer functions (MTFs) that are possibly inherited from these inputs (Gummer et al., 1988a; Huffman & Henson, 1990; Schofield, 2011). MOC neurons also receive ascending projections from cell types in the CN that have wide dynamic ranges (WDR), including T-stellate cells (Blackburn & Sachs, 1990; Guinan, 2018; Romero & Trussell, 2021) and neurons in the small cell cap, which receive low- and medium-spontaneous-rate AN inputs (Ye et al., 2000; Fekete et al., 1984) (Fig. 1). Although some CN neurons have bandpass MTFs, they tend to have broad, shallow peaks and best modulation frequencies (BMFs) that are higher than BMFs for neurons in the IC (Nelson and Carney, 2004; Rhode and Greenberg, 1994). Either source (i.e., IC or CN) of modulation tuning, or even modulation tuning that could arise *de novo* at the level of the MOC, could contribute to the MOC sensitivity to fluctuations. Nevertheless, a parsimonious explanation is that MOC neurons inherit their bandpass MTFs from IC inputs rather than from CN inputs. For the proposed model, IC projections to the MOC provide neural-fluctuation information to the MOC gain-control system, as follows: 1) IC neurons that are excited by fluctuations (band-enhanced (BE) IC cells) effectively reduce cochlear gain in response to fluctuating inputs, 2) this reduction in cochlear gain increases AN fluctuation depth by effectively increasing the IHC dynamic range, 3) this increase in AN fluctuation depth results in a further increase in IC activity until the gain stops decreasing, either because a minimum cochlear gain is reached or when fluctuation-driven feedback is balanced by WDR driven feedback (described below). This fluctuation-driven feedback loop, in which BE IC activity ultimately reinforces BE IC activity, is effectively a positive feedback loop. Note that, a decrease in AN fluctuation depth results in positive feedback that works in the opposite direction, that is, a reduction in IC responses ultimately leads to a further reduction in IC responses.

**FIG. 1.**
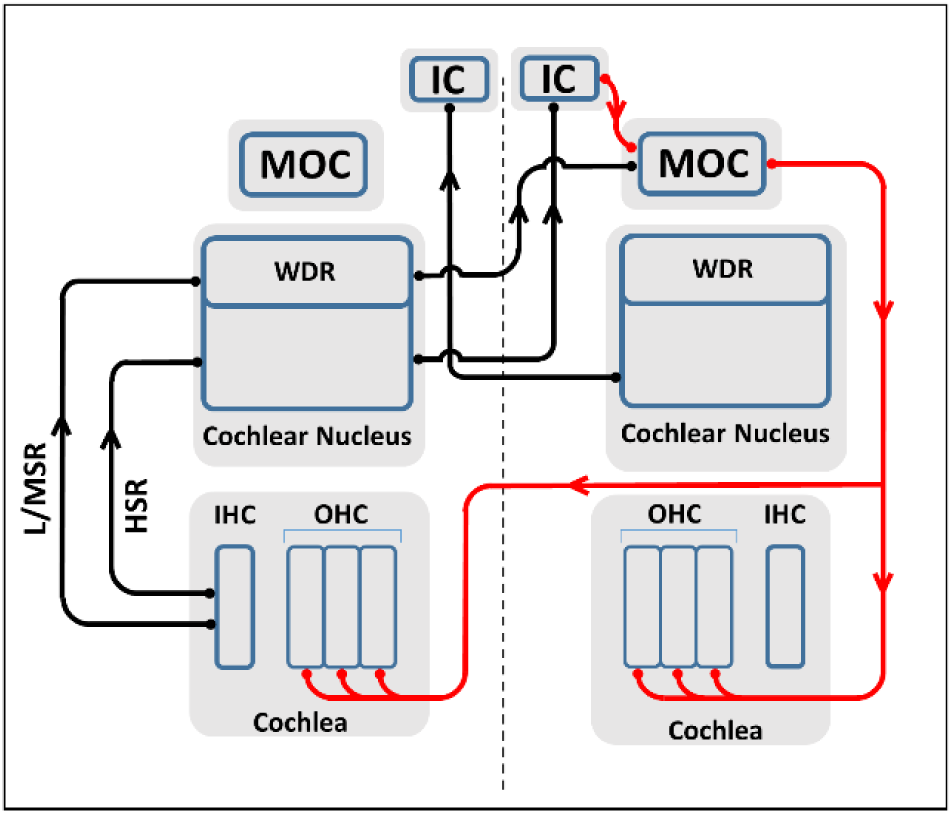
(color online) Schematic illustration of the peripheral afferent (black arrows) and MOC efferent (red arrows) pathways. Here for a simple presentation we have shown the afferent pathways on the left side and the efferent pathways on the right side, but both pathways exist on both sides of the brainstem.

The positive-feedback mechanism described above could contribute to spectral coding of complex sounds, as follows (see Carney, 2018): 1) AN frequency channels tuned near local peaks in the stimulus spectrum have responses that are “captured” by the spectral peak, and thus have relatively small fluctuations (Deng et al., 1987; Delgutte and Kiang, 1984; Young and Sachs, 1979). Consequently, for these frequency channels, BE IC cells would have lower response rates than adjacent channels tuned to spectral troughs. Activity of MOC neurons reduces cochlear gain in a frequency/place-specific manner (Guinan, 2018); therefore, input to the MOC from BE IC channels near a spectral peak would be relatively small, as compared to that from channels within spectral troughs that are driven by large fluctuations. The relatively higher cochlear gain for channels near spectral peaks would drive IHCs closer to saturation, resulting in further reduction in the neural fluctuations in AN HSR responses over time (i.e., positive feedback). 2) For frequency channels tuned away from spectral peaks, AN fluctuations are larger and BE IC cells therefore have higher response rates. The higher rates in the BE IC input to the MOC would result in a reduction in cochlear gain, reducing IHC saturation and further increasing fluctuation amplitudes in the AN response in the frequency channels tuned away from spectral peaks. Overall, the IC input to MOC neurons provides a positive feedback that would enhance the fluctuation contrast across AN channels by decreasing the fluctuating amplitude in frequency channels near spectral peaks and increasing the fluctuations in other channels. This feedback potentially enhances the neural representation of spectral peaks and valleys at the level of the IC (see Figs. 2 and 3 in Carney, 2018).

**FIG. 2.**
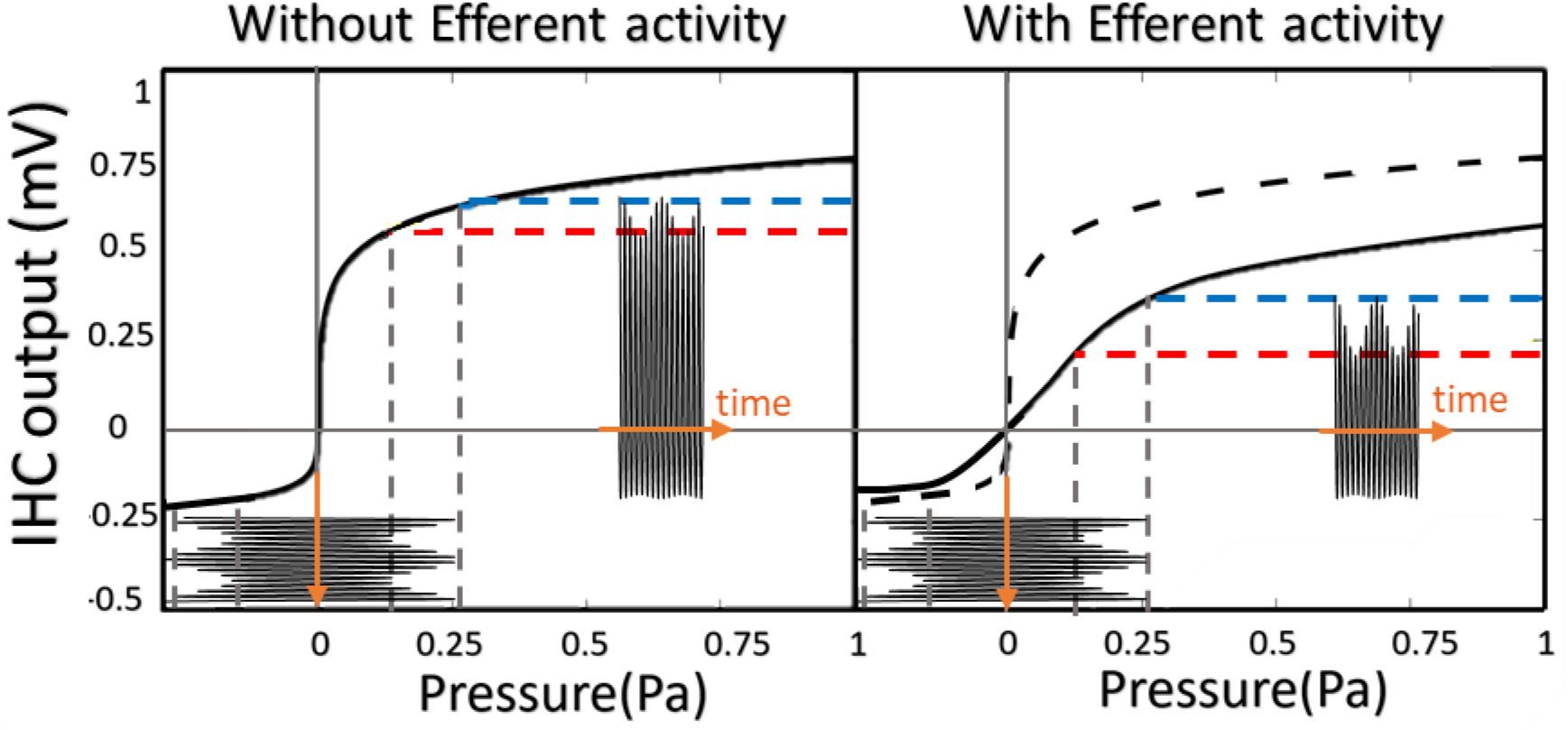
(color online) Effect of MOC efferent activity on IHC input-output function. Left panel: The effective modulation depth in IHC output voltage is limited due to the saturation. Right panel: with the same input pressure, effective modulation depth is increased due to the MOC efferent activity and decrease in gain.

**FIG. 3.**
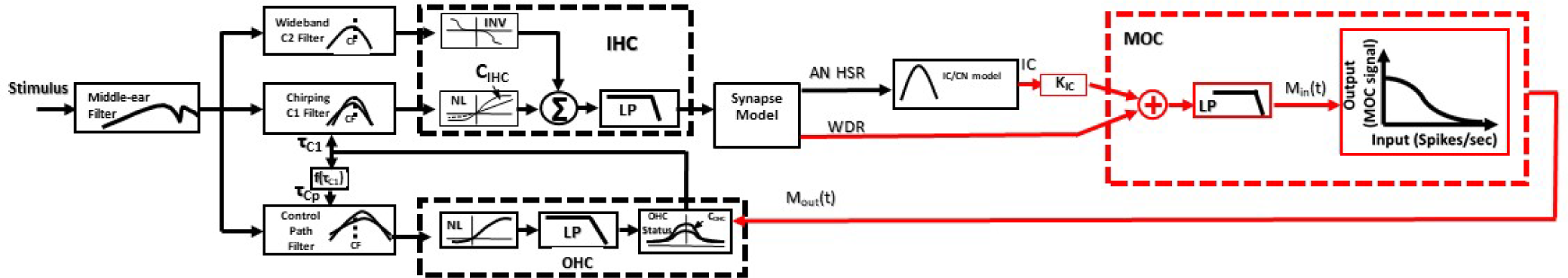
(color online) Schematic illustrating overall structure of the new model components (red) with respect to the Zilany et al. (2014) AN model and Mao et al. (2013) IC/CN model (black) (adapted from Zilany et al., 2014, with permission of Acoustical Society of America.). The Zilany model includes the feedforward blocks simulating middle-ear filter, cochlear filtering (labeled C1, C2, and Control Path Filters) and two stages consisting of filters and nonlinearities for simulation IHC and OHC units (boxes marked by black dashed lines). The new model components include the two feedback signals from IC and WDR cells to the MOC stage (dashed red box), where the two input signals are scaled and summed before going through a first-order low-pass filter. The low-pass filtered signal goes to the rational input-output MOC function, which maps the input signal to a cochlear-gain scaler between 0 and 1, for that time sample.

In the proposed model, the other input to the MOC stage is from the WDR neurons in the CN, which carry the stimulus sound-level information. This level-dependent input to the MOC acts as a negative feedback system, i.e., an increase in stimulus level would increase activity of the WDR neurons which excite MOC neurons, ultimately reducing cochlear gain. This negative feedback is hypothesized to keep the operating point of the saturating IHC transducer function near its knee point, to ensure sufficient baseline fluctuation contrast across frequency channels. The IC feedback signal, described above, would then enhance this baseline contrast. If there were no contribution to gain control from the WDR cells, the AN channels could all have fluctuating or flat responses when exposed to stimuli with low or high levels, respectively, regardless of the peaks and valleys in the stimulus spectrum. Both extremes would reduce the contrast in fluctuation depth across IC frequency channels.

According to the neural-fluctuation profile framework and the feedback mechanisms of the MOC efferent system described above, sustained fluctuations in a stimulus could cause a progressive decrease in cochlear gain over time. As a result, in response to amplitude-modulated (AM) noise, the average rate of BE IC neurons is expected to increase over a time course of 100-200 ms due to the MOC-related reduction in cochlear gain. The rationale for this prediction is that the low-frequency fluctuations of AN fiber responses to modulated noise stimuli excite IC neurons with BE MTFs. BE IC neurons are assumed to excite MOC cells, and MOC activity decreases cochlear gain. Reduction in cochlear gain effectively decreases the slope of the IHC transduction nonlinearity and therefore increases the effective modulation depth at the output of the IHC (Fig. 2), ultimately increasing the BE IC response. Thus, IC rates would increase over time until the efferent control system is stabilized. Interestingly, re-examination of IC responses to AM noise in awake rabbits (Carney et al., 2014) supported this hypothesis and revealed rates that increased over a time course that is consistent with the dynamics of MOC efferent gain control (Farhadi et al., 2021). This increase in IC rate is an unexpected effect in neural responses, which typically adapt and therefore exhibit a decrease, rather than increase, in the neural rate over time. We quantified the increase in IC average rate over the time course of responses to AM noise and used that information to adjust the parameters of the MOC efferent model proposed in this work.

Here, we refine and test an AN and subcortical model with both fluctuation- and WDR-driven gain control (Farhadi et al., 2021). We first introduce the novel control system used to implement the MOC efferent pathway. Next, we describe how the parameters of the MOC efferent model were adjusted based on a physiological dataset from a previous study of IC responses to AM noise from awake rabbits (Carney et al., 2014). We then show that the model with efferents outperforms a model without efferents in simulating the dynamics of the IC response to AM noise stimuli. This finding supports the hypothesis that the MOC efferent system is a potential mechanism underlying the increase in rate of IC cells over the time course of AM noise. Implications of the proposed model for responses to other sounds, and interactions with other neural mechanisms involved in long-term changes in response rates, will be discussed.

## II. METHODS

We adapted existing models that were designed to simulate responses of AN fibers (Zilany et al., 2009, 2014) and IC cells (Mao et al., 2013) to implement the proposed model with MOC efferent gain control. First, we briefly introduce the original AN and IC models. Then, we describe the modifications that were applied, including changes to the structure of the existing models and new extensions added to simulate the MOC efferent system. Finally, we introduce the methods used to estimate the MOC efferent-system model parameters.

### A. Existing models

For the original Zilany et al. (2009, 2014) AN model, the stimulus first passes through a filter that simulates the middle-ear transfer function. The output of the middle-ear filter is the input to the cochlear-model stage (Fig. 3). Cochlear tuning is modeled using two filters, a relatively sharp chirping filter in the signal path tuned at the characteristic frequency (CF) and a wider bandpass filter in the feedforward control path tuned slightly higher than CF (Zhang et al., 2001; Tan & Carney, 2003; Zilany & Bruce, 2006). The output of the control-path filter and following OHC model stage adjusts the time constant (and thus the gain and bandwidth) of the chirping filter. The cochlear-gain parameter (C_OHC_) in the original model is set to one to simulate normal hearing. The IHC is modeled with a nonlinear transduction function followed by a low-pass filter. IHC sensitivity (C_IHC_) is set to 1 to simulate normal-hearing IHC function. The AN model is based on human Q10 values (Shera et al., 2002).

We used the BE IC model from Mao et al. (2013), which consists of a band-pass filter centered at the BMF of the IC cell. The response of a simulated HSR AN model was the input to the IC model; this structure is a simplified version of an IC model for AM tuning that also included a CN stage (Nelson and Carney, 2004).

### B. Proposed model with MOC efferents

One of the challenges of including the efferent system in the computational AN model was to reconfigure the implementation of the model to work sample by sample. In previous IC model implementations (e.g., Mao et al., 2013), the AN model output for the entire stimulus waveform served as the input to the IC stage. In the sample-by-sample configuration of the proposed efferent model (Fig. 3), a new value of C_OHC_(t) was applied to each sample to adjust cochlear gain based on the output of a new stage: the model MOC (Fig. 3, dashed red box). The dynamics of the MOC system were modeled with a first-order low-pass filter, thus the output of the MOC model was a function of the current and previous time samples of the MOC input signal, which was a combination of the IC-model rate function (time-varying rate, in spikes/sec) and the CN WDR response provided by a LSR AN-model rate-function. For simplicity, we used the LSR AN model response to represent all WDR inputs to the MOC, such as the T-stellate cells in the anteroventral cochlear nucleus (Oertel et al., 2011; Romero & Trussell, 2021) and cells in the small cell cap of the CN that receive exclusive inputs from LSR and MSR AN fibers (Fekete et al., 1984; Ryugo, 2008). In the sample-by-sample implementation, the previous states of all the filters of the AN model were preserved in order to process each time sample of the simulated model response.

In the proposed model, the output of the MOC efferent stage controlled the gain parameter based on the current and previous samples of the neural signals at the input to this stage. Specifically, the output of the MOC stage, a scaler between 0 to 1, was multiplied by the OHC gain parameter (C_OHC_) to modulate the cochlear gain. This multiplication occurred for every time sample. The model with this new sample-by-sample configuration was tested to ensure that it matched the response of the original model when the efferent stage was deactivated.

The two inputs to the proposed MOC model (Fig. 3, red) were the descending IC feedback signal and the ascending WDR signal, which were combined to determine the degree of cochlear gain reduction. Specifically, the excitatory input to the MOC stage was the sum of the IC and the WDR rate functions. A constant scalar, K_IC_, was applied to the IC rate signal to approximately balance the amplitudes of the two inputs to the MOC. The AN model has higher average rates than the IC model, and combining these signals without scaling would result in an input signal to MOC that would not be not sensitive to changes in the IC model response. The MOC system was modeled with a low-pass filter representing dynamics of the MOC efferent system, followed by a decaying rational function (Eqn. 1) that mapped the MOC inputs to an output signal that modulated the cochlear-gain parameter, C_OHC_, on a sample-by-sample basis. The rational function, chosen to be second-order to ensure asymptotic behavior for both small and large input signals, was described by

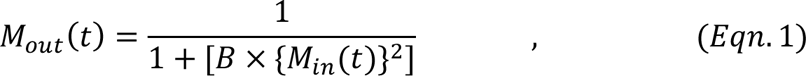

where *M_in_*(*t*) is the sum of the responses from the WDR and IC paths to the MOC unit after low-pass filtering, and B is a constant that adjusts the shape of MOC input-output function (Fig. 3). The optimization of the value of parameter B is discussed later (Section C of the Methods).

Figure 4 (upper panel) shows the intermediate responses at each level of the composite model to a tone-in-noise stimulus. The first 200 ms of the stimulus is noise only (Fig. 4, black). The BE IC response to this initial noise is robust (Fig. 4, purple); because of the fluctuation in AN HSR rates (Fig. 4, blue), thus driving the MOC model and decreasing the cochlear gain (Fig. 4, green). During the first 200 ms the gain decreases over time and as a result the fluctuations in the AN HSR response increase over time (Fig. 4, blue-see also Fig 4A-D for zoomed-in views of AN HSR responses corresponding to labelled portions in Fig. 4, upper panel). The increase in depth of fluctuations in the AN HSR response over time (Figs. 4C, D) resulted in an effective decrease in cochlear gain from the MOC stage and subsequent increase in IC rate over time (Fig. 4, purple). Between 200 and 500 ms, a tone at the model CF (4 kHz) was added to the noise, resulting in saturation of the IHC (not shown) and flattening the neural fluctuations in the AN HSR response (Fig. 4, blue). As a result of the reduced AN HSR neural fluctuation depths, the BE IC rate response decreased. The reduced IC rate resulted in a weaker MOC input and thus a relative increase in cochlear gain (Fig. 4, green). Note that the mean rate of the AN HSR fiber was not changed appreciably by the addition of the tone, due to rate saturation; however, the model WDR rate function increased during presentation of the tone. This increase in rate of the WDR response over the time course of the tone occurred because the average rate was not saturated for WDR neurons. After the tone ended, the IHC was no longer saturated, AN fluctuations reappeared, and the IC response increased. The increase in the IC response excited the MOC stage, which resulted in a reduction in cochlear gain. The changes in the WDR response rate (Fig.4, red) also influenced cochlear gain. As described above, the WDR signal maintains the IHC operating point such that the IC response has an effect on increasing spectral contrast via changes in cochlear gain. In contrast to the CF channel tuned to the tone frequency (Fig. 4), AN fibers tuned away from the tone would have responses that were dominated by the noise masker; at moderate sound levels, these responses would have strong fluctuations. In summary, the introduction of a tone in an ongoing noise results in a flat AN response that is temporally flanked by fluctuating responses and thereby may provide a cue for detecting a tone in noise. The responses of the model without efferent gain control did not show changes over time during each portion of the stimulus, as expected (Fig.4, upper panel left, and Figs. 4A, B).

**FIG. 4.**
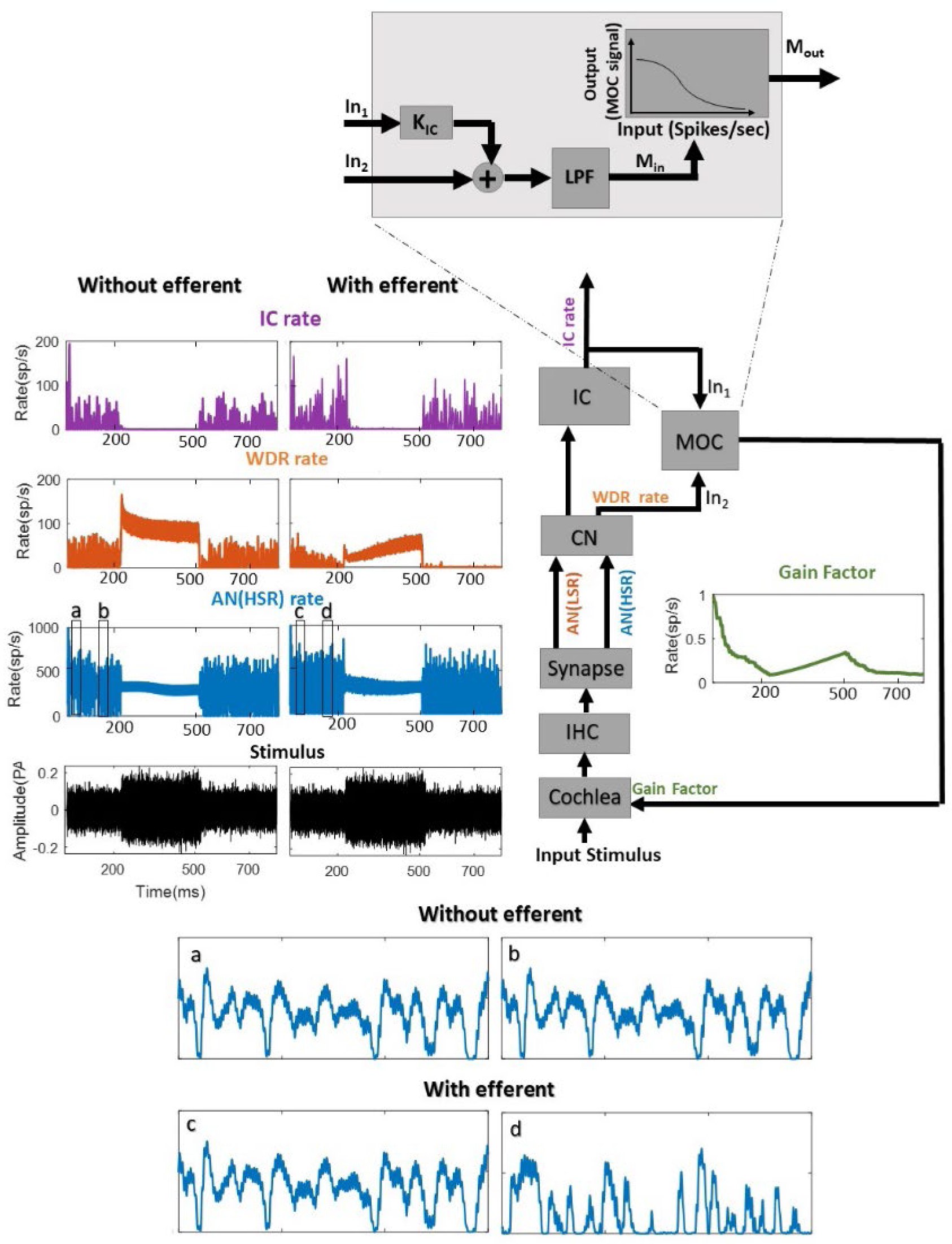
(color online) The MOC efferent model diagram with intermediate responses at different levels of auditory periphery in response to tone-in-noise stimulus. The stimulus (black) is initially a wide band noise; after 200 ms a tone is added to the noise for a duration of 300 ms. The response of the model AN (HSR) stage (blue) shows that during the noise-only stimulus (first 200 ms) the fluctuation depth increases over time as the gain decreases (green), and as a result the IC (purple) rate increases. After the addition of the tone (200-500 ms) the fluctuation depth in the AN HSR response is reduced, resulting in a drop in IC response rate. As the In1 input to the MOC block (on the right side of this figure) decreases, the gain starts to recover (green). After the offset of the tone (500-700 ms) the gain decreases again, and the AN fluctuations and IC rate increase.

FIG. 4. (color online) The MOC efferent model diagram with intermediate responses at different levels of auditory periphery in e to tone-in-noise stimulus. The stimulus (black) is initially a wide band noise; after 200 ms a tone is added to the noise ration of 300 ms. The response of the model AN (HSR) stage (blue) shows that during the noise-only stimulus (first 200 fluctuation depth increases over time as the gain decreases (green), and as a result the IC (purple) rate increases. After the of the tone (200-500 ms) the fluctuation depth in the AN HSR response is reduced, resulting in a drop in IC response the In1 input to the MOC block (on the right side of this figure) decreases, the gain starts to recover (green). After the offfset the tone (500-700 ms) the gain decreases again, and the AN fluctuations and IC rate increase.

### C. Adjusting the parameters of the MOC efferent model

Parameters of the MOC efferent model include those of the MOC input/output function, *B* (Eqn.1, Fig. 4), the scaler constant for IC rate (*K_IC_*), and the low-pass filter (LPF) time constant (cut-off frequency). We adjusted these parameters based on an existing physiological dataset that contained IC responses in awake rabbits to AM wideband noise (Carney et al., 2014).

#### Stimuli

The stimulus for both the physiological dataset and the model simulation during the parameter fits was a sinusoidally amplitude-modulated (SAM) wideband noise (100 Hz to 20 kHz) with modulation depth that varied from -30 to 0 dB (fully modulated) in 5 dB steps, and an unmodulated wide-band noise with the same bandwidth. The modulation frequency was matched to the best modulation frequency (BMF) of the recorded cells, which ranged from 50 to 100 Hz. The stimulus spectrum level was 36 dB re 20µPa (RMS: 76 dB SPL overall level). The duration of the stimuli was 500 ms, including 50-ms raised-cosine ramps.

#### Physiological data and analysis

The physiological dataset included previously published extracellular, tetrode recordings of responses in the central nucleus of the IC in three awake rabbits. Stimuli were presented to the contralateral ear. Details of the physiological methods can be found in Carney et al. (2014). Briefly, IC cells were classified based on their MTFs into BE, band-suppressed (BS), hybrid, flat, and unusual types (Kim et al., 2015, 2020) (Fig. 5). BE IC cells are excited over a band of modulation frequencies, whereas IC BS cells are suppressed over a band of modulation frequencies (Kim et al., 2020). Hybrid cells have rates that are enhanced over one band of modulation frequencies, and suppressed over another range (Kim et al., 2020). The focus of this study was on BE cells because this IC MTF type was hypothesized by Gummer et al., (1988b) to project to MOC neurons, based on the BE MTFs reported for MOC neurons. MTF shapes were classified similar to Kim et al. (2020), using a criterion ratio of 1.2 peak MTF rate to the reference rate response to an unmodulated wideband noise which yielded approximately equal numbers of BE and BS MTFs (Fig. 5). The frequency of the peak rate in the MTF identifies the cell’s BMF. The physiological responses used in this study had stimulus modulation frequencies that were within an octave of the recorded cell’s BMF.

**FIG. 5.**
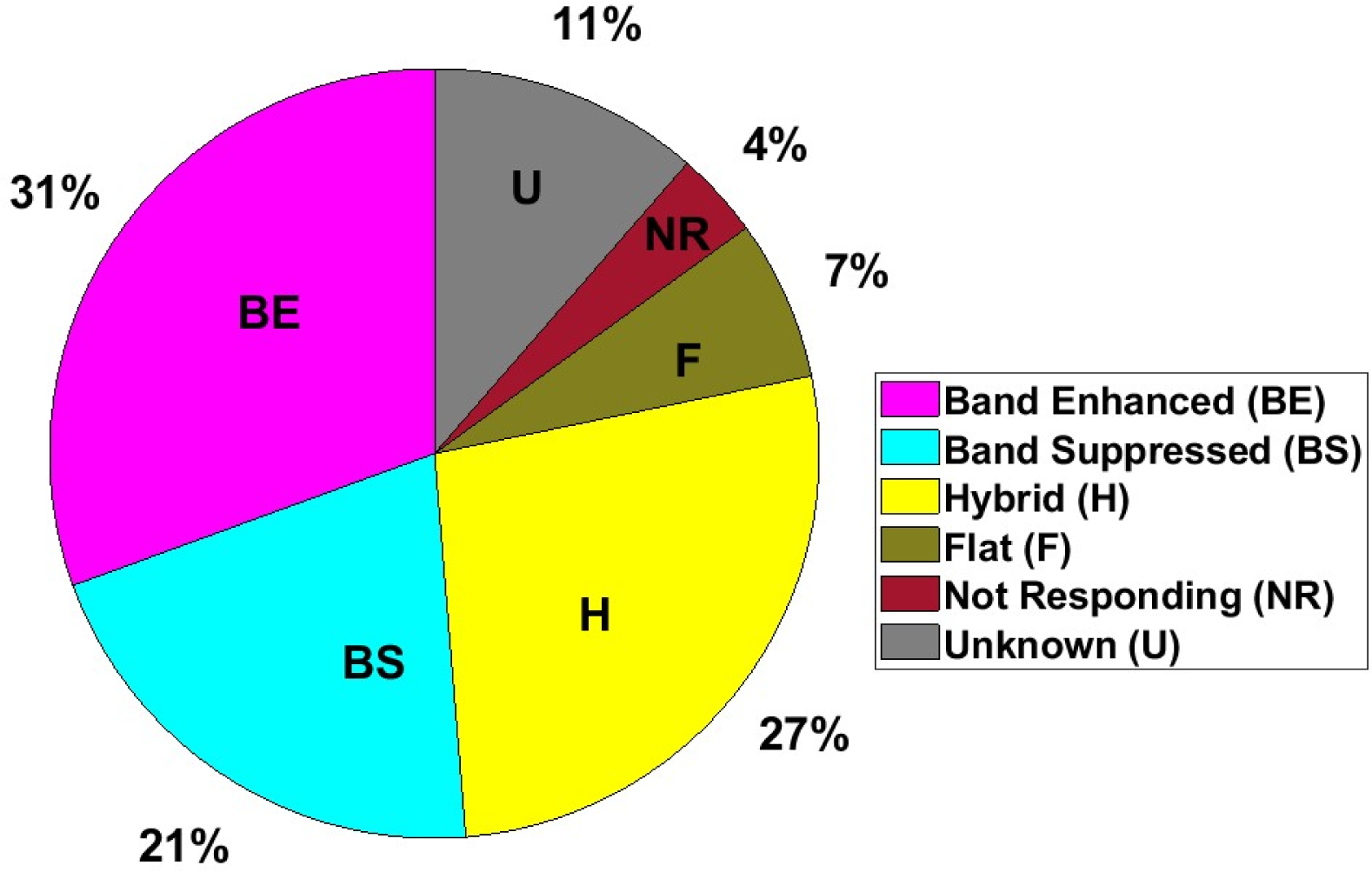
(color online) Distribution of MTF classes of the IC neurons (N=193) in the physiological dataset (Carney et al., 2014) using the MTF-categorization method of Kim et al. (2020) (D. O. Kim et al., 2020)

To analyze the physiological responses to AM noise, a Gaussian filter was used to smooth the spike trains to avoid aliasing introduced by PSTH binwidth (Lehky, 2010). The standard deviation of the Gaussian filter was inversely proportional to each cell’s BMF; the number of points in the filter was six times the standard deviation of the Gaussian window. Responses were averaged over 50 repetitions of the stimulus.

Trends in firing rate over the course of the response, after an initial 200 ms to exclude the onset response, were characterized using a simple linear regression. Based on the efferent model framework, the BE IC rates were hypothesized to increase over time as a result of reduction in cochlear gain and increase in the temporal fluctuation depths in AN responses. The sign of the linear model’s slope indicated whether the response increased or decreased over time. BE IC responses with rates that increased over time were used to find the parameters in the MOC efferent model by maximizing the correlation between the BE IC spiking rate and the model responses to a stimulus with a modulation depth of -10 dB. A modulation depth of -10 dB was chosen here because the majority of the IC neurons had strong increases in rate over time at this modulation depth as compared to higher modulation depths, for which IC rates sometimes saturated. A brute-force method was used to search a reasonable range (*K_IC_* between 1 to 50, *B* between 0.001 to 0.1, and *τ_MOC_* between 10 ms to 500 ms) of parameter values and find a set of values that maximized the correlation between the model response and the physiological data over the analysis window. Parameters were estimated for each neuron in the dataset. We simultaneously fit the time constant, *τ_MOC_*, the rational function parameter, *B* (Eqn. 1), and the IC constant scaler, *K_IC_*. A 300-ms-duration analysis window was used for both physiological and model responses. The analysis window for model responses started at 200 ms in order to exclude the onset response, as was done for the physiological responses, and continued until the end of stimulus. Properties of the model such as CF, BMF, and modulation frequency were matched to each cell in the physiological dataset. Based on the parameter value fits for each of the 33 neurons in the dataset, a distribution of the parameters was established, and the medians of the distributions were chosen for the final model parameter values. The sensitivity of the model to the parameter values was studied by comparing the average correlation between the physiological and the model responses when each model parameter was either set to the value fit to each neuron or to the median parameter value for the population.

## III. RESULTS

Section A below reviews our hypothesis regarding the MOC efferent model response to AM noise stimuli, analysis of the physiological dataset in response to AM noise, and implementation of the MOC efferent model with its parameters (*K_IC_, B,* and *τ_MOC_*). Section B describes the distribution of the optimum parameters in the dataset and compares the responses of models with and without the MOC efferent control system to physiological data. In section C we use the new model with MOC efferent control to simulate responses to a few different stimuli and compare these responses with physiological data from the literature.

### A. The physiological dataset

Consistent with our hypothesis, the IC rates increased over time in most IC neurons with BE MTFs. Figure 6 presents six examples of BE IC neuron responses that illustrate the trend of increasing rate over time; these examples were chosen as they have different BMFs and CFs. The left panel in each example in Fig. 6 is the rate MTF, and the right panel is the IC response to AM noise modulated at a frequency near the neuron’s BMF, with a range of modulation depths (different colored lines). The gray time range is the time window after the onset response over which we analyzed the increase in rate. All of the examples in Fig. 6 had increasing rate over time during the analysis window, especially in response to lower modulation depths. In some neurons (e.g., Fig. 6B) the rate was nearly constant over time in response to fully modulated noise, but the IC rate was high for this modulation depth from the beginning of the analysis window. In the framework of our hypothesis, this observation suggests that because of the strong modulation in the stimulus, the fluctuation in AN PSTHs was large enough to maximize the IC rate, such that a change in cochlear gain as a result of MOC efferent activity could not further increase the IC rate. In these examples, the PSTHs in response to the unmodulated noise stimulus also showed an increasing trend in rate. This trend in the noise responses could also be explained by efferent-system activity, because unmodulated wideband noise includes inherent temporal fluctuations after peripheral filtering, which would increase IC rate and lower the cochlear gain.

**FIG. 6.**
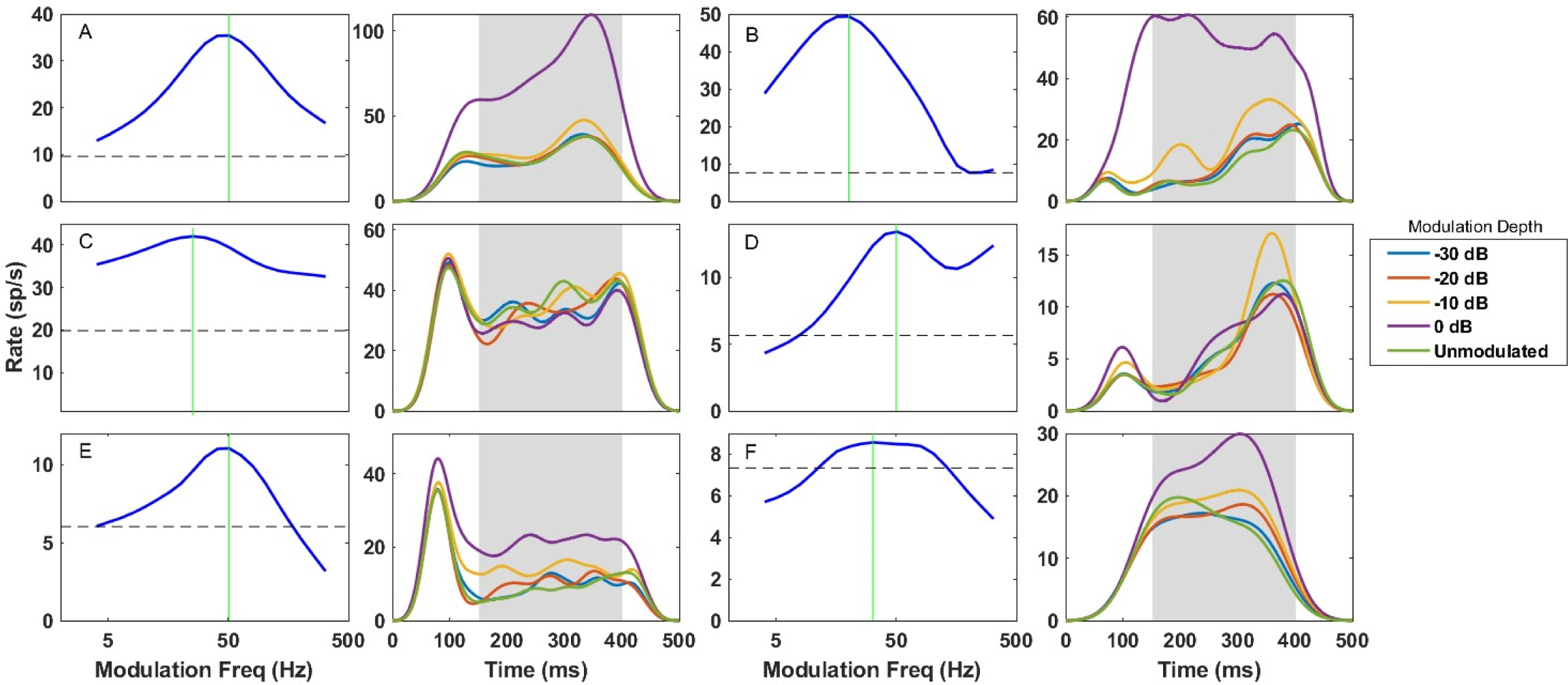
(color online) A-F) Six examples of BE IC neuron smoothed responses to AM noise stimulus. The left-hand panel in each example is the MTF for AM noise stimuli; the vertical green lines indicate BMF. The right-hand panels show smoothed PSTHs that illustrate the time course of the rate during responses to AM noise stimuli with a modulation frequency near each neuron’s BMF and a range of modulation depths (see legend).

The trend of increasing rate over time (See Methods Section C) occurred in the majority of the neurons with BE MTFs in response to modulation depths less than 0 dB (i.e. most symbols in Fig. 7 fall above the horizontal line at 0). The percentage of neurons with positive slopes decreased at high modulation depths (see Fig. 6). As described above, the decrease at high modulation depths can be explained by a ceiling effect on the IC rate. In general, neural response rates are typically expected to decrease over time during sustained stimuli, as a result of rate adaptation, making this increase in rate particularly striking. The finding that that majority of the BE IC cells had rates that increased over the time course of the AM noise stimuli is consistent with the hypothesis that MOC efferent activity results in an increase in BE IC rates over time.

**FIG. 7.**
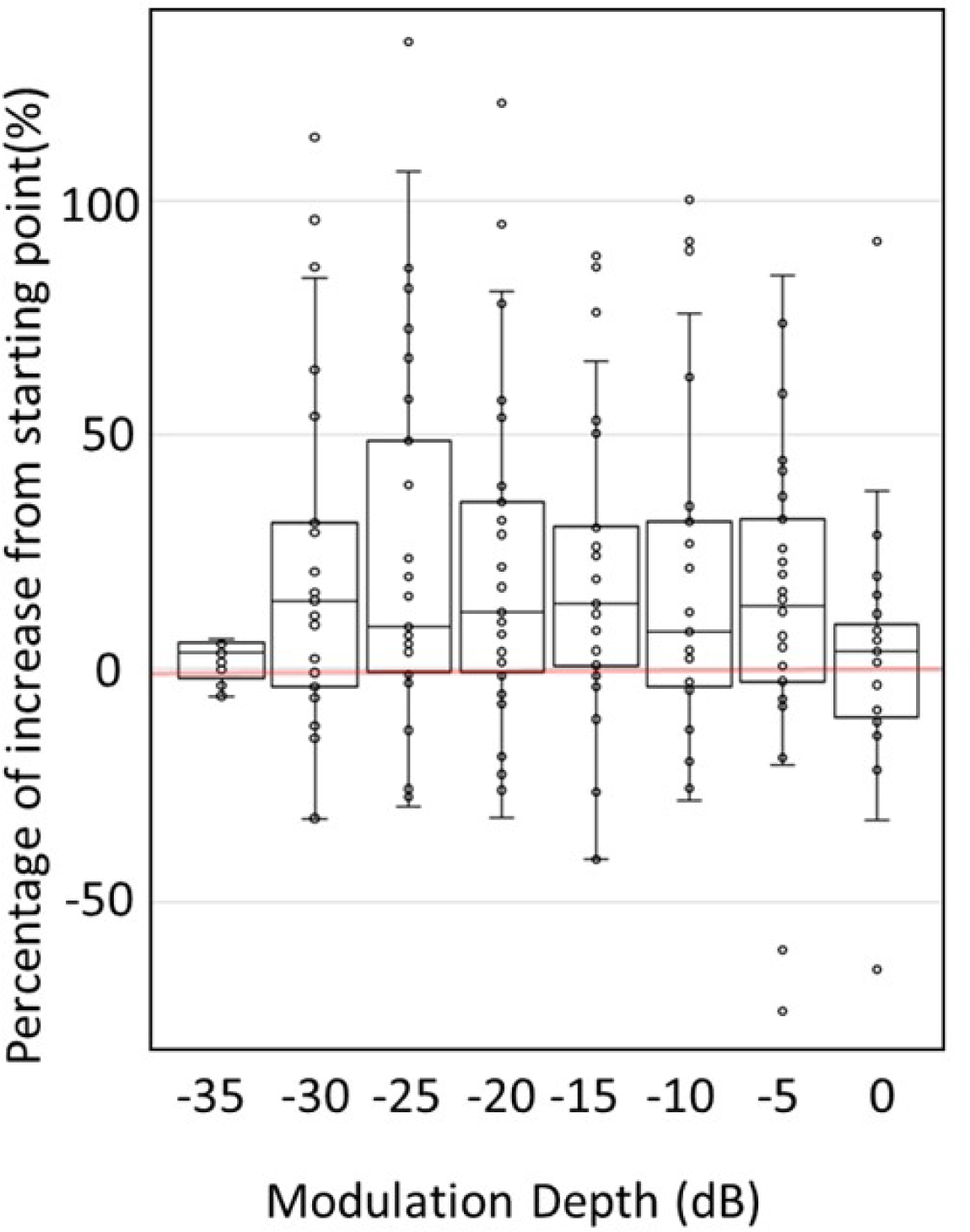
Percentage increase in rate of BE IC neurons, referenced to the rate at the end of the onset response. The majority of 35 studied BE neurons had rates that increased over the steady-state time course of wideband AM noise stimuli.

### B. Adjusting model parameters

The distributions of the optimized parameters for neurons in this study were broad (Fig.8). The median of the distribution for each parameter was used as the final value for that parameter (K_IC_=20, τ_MOC_=260 ms, B=0.01) (Eqn.1). The median of the estimated time constant distribution was 260 ms, which was used to determine the cutoff frequency of the LPF in the MOC model, τ_MOC_. This value is consistent with values of the time constant that have been suggested for the MOC efferent system based on other types of physiological and psychophysical measurements (Backus & Guinan, 2006; Roverud & Strickland, 2010; Warren & Liberman, 1989). The median of the distribution of correlations between the model responses and each of the IC neuron responses in the physiological dataset was 0.025 for models without efferent feedback. These correlations illustrate that the model without efferents did not simulate the increasing trend in rate over time in the physiological dataset. The response of the model without efferents had a decrease in rate consistent with neural adaptation that arises in the AN model responses (Zilany et al., 2014). Simulations using a MOC efferent model with parameters identified for each neuron resulted in a correlation of 0.89 between the data and model responses. The correlation reduced to 0.43 after replacing the individual values with the median values of the distributions estimated for each parameter. Some of the correlations in this condition were negative, consistent with the wide distribution of the optimum parameters for each neuron. We performed cross-validation analysis with 30 validation neurons and 5 test neurons, using a bootstrap analysis and 100 repetitions with replacement. The median of the test correlation was 0.2 while the validation correlation was 0.43. This result showed that the model was sensitive to the parameter values, but still simulated the increased rate in response to AM noise using the median parameter values. To test the chosen values for the model parameters, we show the median correlations between the physiology and the model responses for the values in the optimization search window and evaluate the effect of each of these parameters on this correlation (Fig. 9A). The final values for the model parameter (*K_IC_* = 20, τ_MOC_ = 260 ms, B = 0.01) had the highest correlation value points on the surface plots in Fig. 9, supporting the validation of these values.

**FIG. 8.**
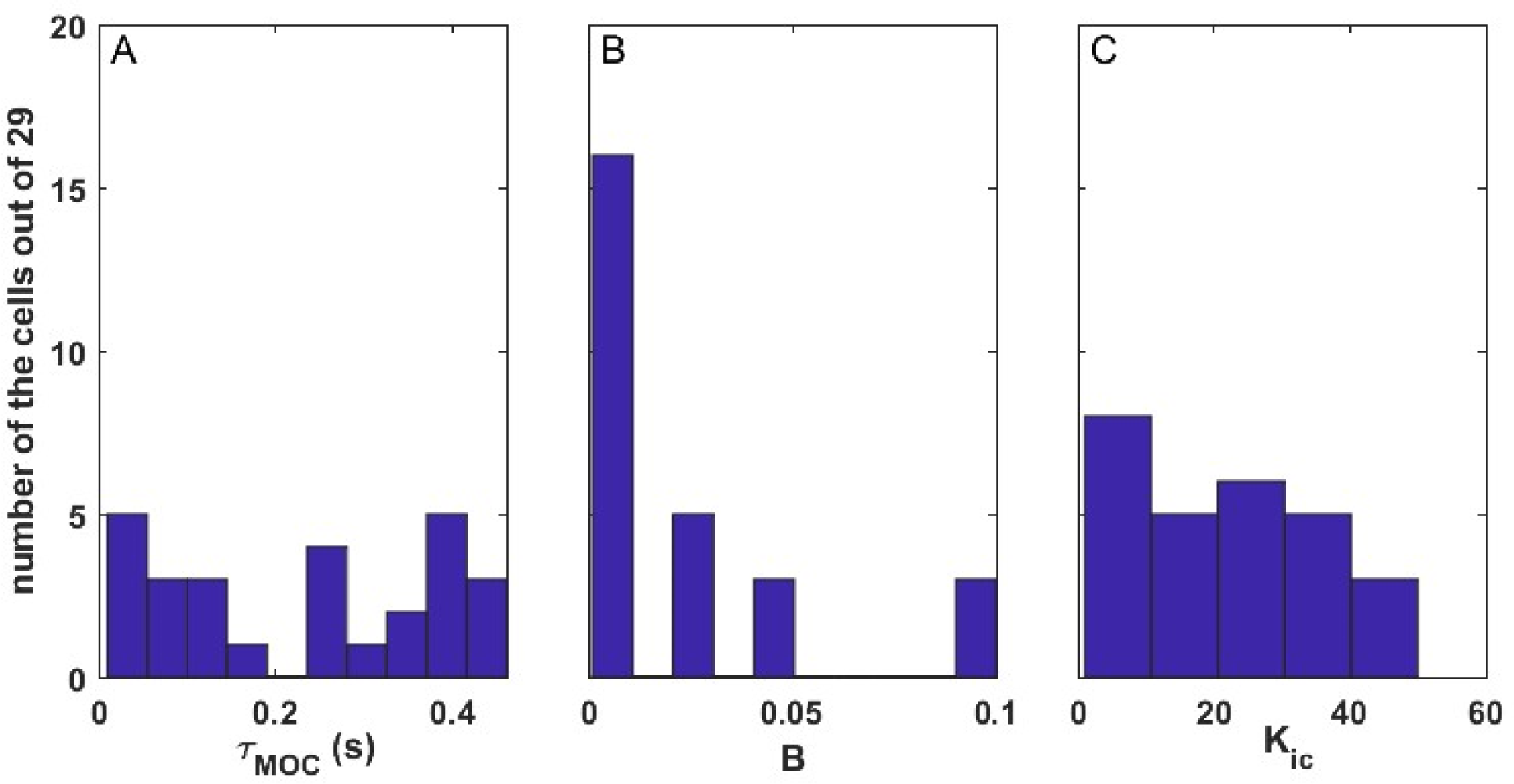
(color online) Distribution of the fit values of, A) time constant (τ_MOC_), B) the rational function parameter (B), and C) the IC scaler (*K_IC_*) for the model for 29 neurons in this study for which the rates increased during the SAM stimuli, the MTF type was BE, and the modulating frequency of the stim-ulus matched the BMF of the cell.

**FIG. 9.**
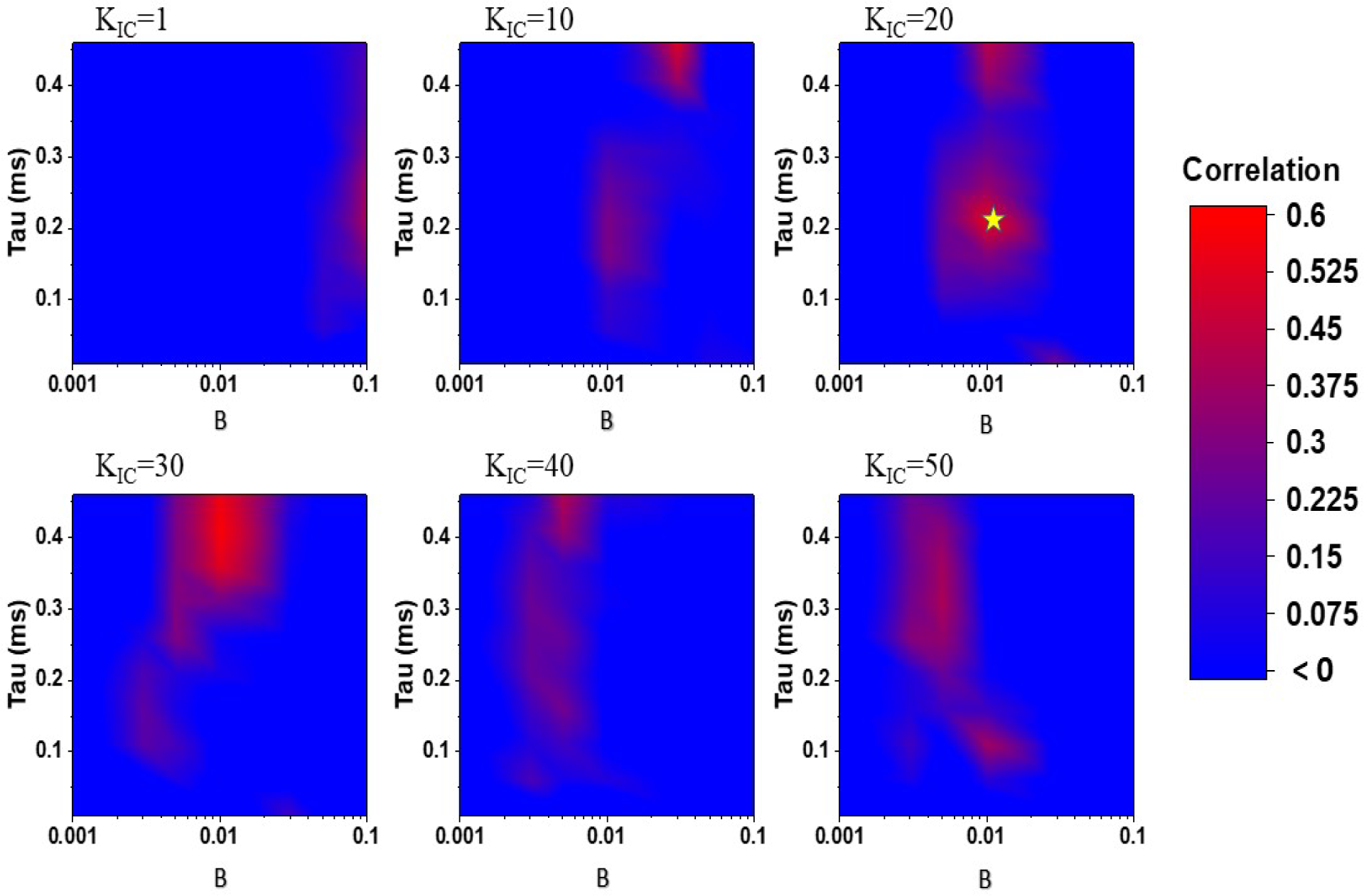
(color online) Correlations between the IC responses and model simulations for various model parameter combinations. Each panel shows the correlation with the changing parameters ‘B’ and ‘τ’ while K_IC_ is held constant at the values displayed at the panel title. All correlations below zero are shown as blue.

For a better understanding of the effect of each of the parameters on the model IC response to AM noise stimuli, Fig. 10 illustrates model responses over time for different combinations of parameter values. Each panel shows responses for the model with the value of one parameter set to five values within the optimization search window, while keeping the other two parameters set to the median of the distributions in Fig.8. For instance, in Fig. 10A, the parameters *B* and *K_IC_* were set to 0.01 and 20, respectively, while the time constant, τ_MOC_, was varied between 10 to 460 ms. For lower values of the time constant, the increase in rate over time evident in the physiological dataset, was not present in the model response. When the time constant was increased to 260 ms - the mean of the distribution in Fig. 8 - an increase in rate over time was observed for the model responses. Further increase in the value of the time constant resulted in a decrease in the slope of the rate increase. Similarly, model responses for a range of values of the parameters *B* (Fig. 10B) and *K_IC_* (Fig. 10C) illustrate how the change in response rate over time was influenced by a range of parameter values. These responses illustrate the sensitivity of the model to the parameter values, and further validate the parameter values chosen to match the physiological responses. Figure 11 shows responses to AM noise for three representative BE IC cells (left column), for models without MOC efferents (middle column), and for the model with efferents, using the median values for the model parameters (right column). The models had BMFs and CFs that matched the example neurons. The model with MOC efferent feedback simulated the increase in rate over time, consistent with the hypothesis that a model with the MOC efferent system could simulate the response to AM noise more accurately than the model without efferents.

**FIG. 10.**
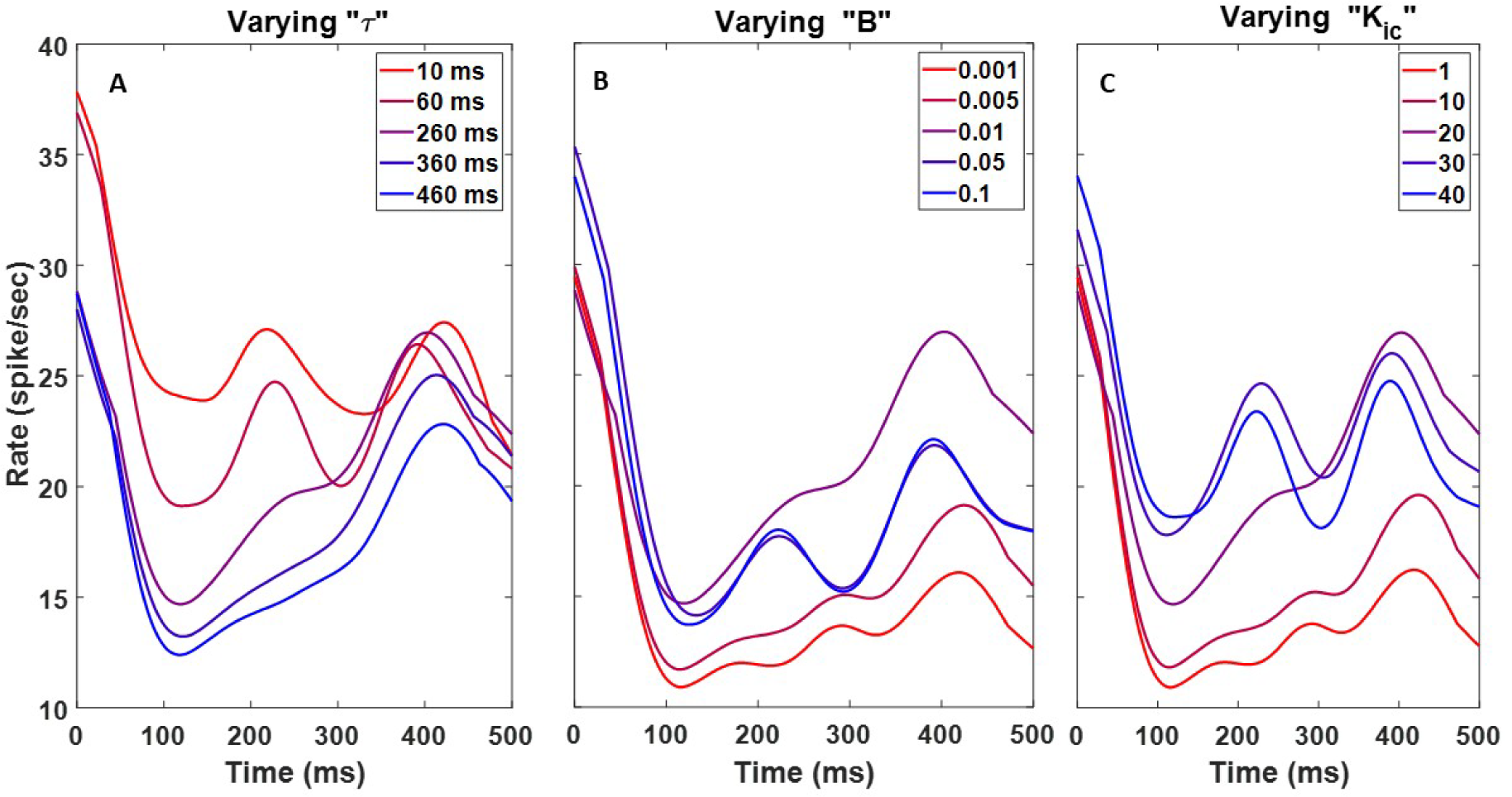
(color online) Effect of changing each of the efferent model parameters on IC model responses to AM noise stimuli over time. A) the time constant parameter (τ_MOC_) was varied, while the other two parameters (*B* and *K_IC_*) were fixed to the median values of the best-fit parameter-value distributions (Fig. 8B. B, C) Model responses for five values of the parameters *B* and *K_IC_*, respectively.

**FIG. 11.**
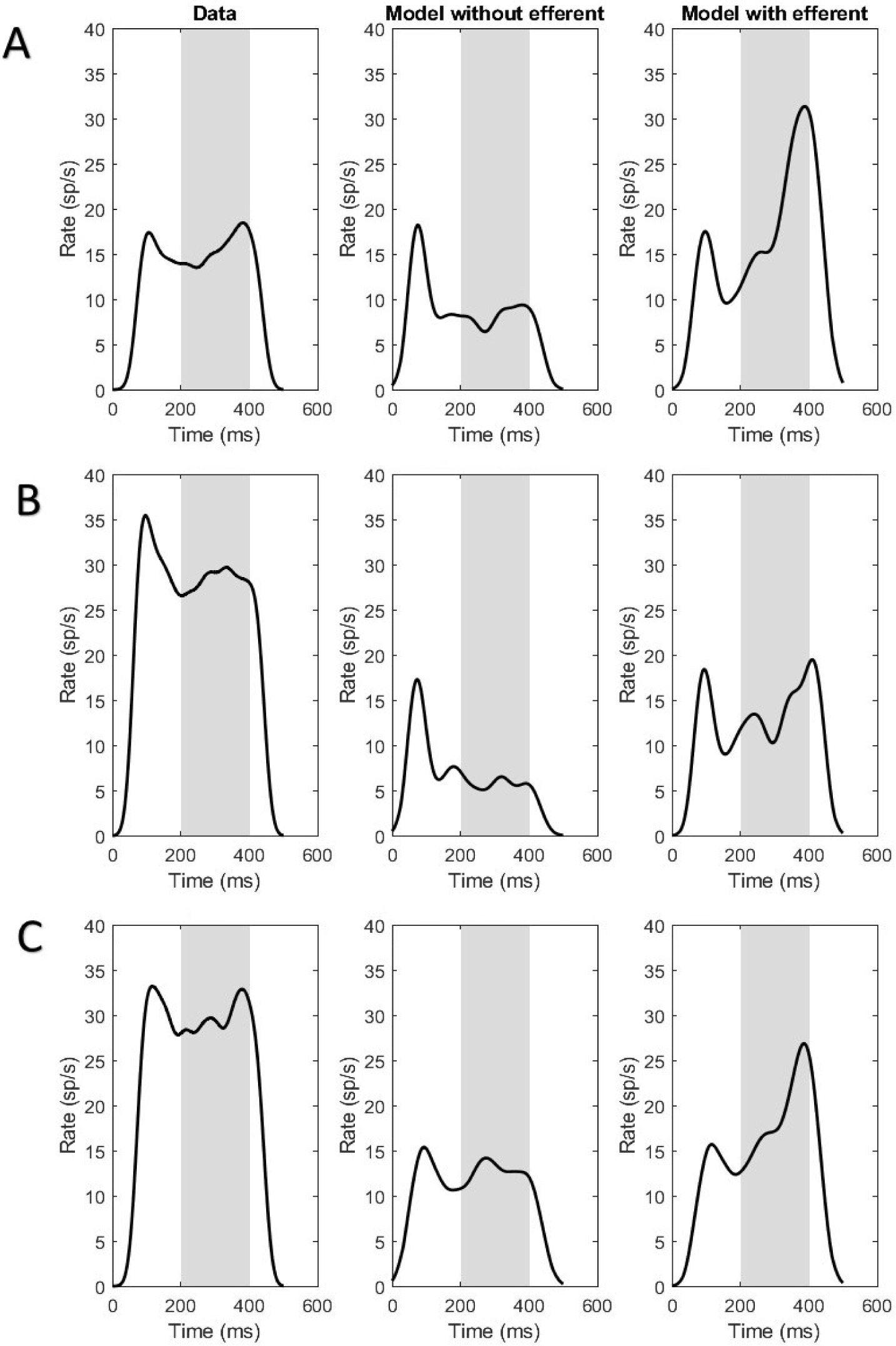
A-C) Examples of physiological data from three BE IC cells in awake rabbit (left panels), with simulated IC responses to AM noise with modulation depth of -10 dB without (center) and with (right) efferent activation. The CF and BMF of each model neuron were matched to the example neuron. Median values (*K_IC_*=30, τ_MOC_=260 ms, and *B*=0.01) for the model parameters were used. The CFs for these cells were 3.5, 4, 2 kHz, respectively, and the BMFs were all approximately 80 Hz.

### C. Validating the proposed model responses

Model responses to tone stimuli were compared to responses of the previous AN model to verify that introducing the efferent gain-control pathways did not adversely affect the model’s ability to predict AN responses to basic stimuli. It is important to note that the physiological data used for comparison were recorded from ANs of anesthetized animals, in which the MOC efferent system was likely suppressed (Guitton et al., 2004). Thus, complete agreement between the physiological responses of AN fibers in anesthetized animals and the responses of the AN stage of the model with MOC efferents was not expected, especially for complex stimuli (such as AM tones). However, for pure-tone stimuli, Rhode and Kettner (1987) and May and Sachs (1992) showed that anesthesia does not significantly affect post-stimulus time histograms (PSTHs) or rate-level functions of primary-like neurons in the cochlear nucleus (and one presumed AN fiber). Therefore, we validated the proposed model by comparing simulated responses to some baseline stimuli in physiological examples.

#### Model Response to pure tones and recovery of spontaneous rate

Figure 12 shows data and simulated responses to a 500-ms tone at CF followed by 500-ms of silence. The purpose of this task was to study both the tone response and the AN model’s recovery to spontaneous rate. Recovery of spontaneous activity depends on stimulus level and on the spontaneous rate of the fiber (Kiang et al., 1965). Here we compare model responses to physiological recordings from two HSR fibers with CFs of 1.82 and 10.34 kHz from Kiang et al. (1965) (Fig. 12). The model response during the tone is comparable to the physiological examples, and the model shows the same initial pause and time course of recovery after the stimulus as in the physiological responses. Note that the response to a pure tone would primarily activate the CN input to the MOC efferent pathway. The IC stage of the model would mainly respond at the onset and offset of the tone, when the onset/offset ramps form a fluctuation in the stimulus envelope and therefore elicit a response from the fluctuation-sensitive IC neurons (Fig. 12.G).

**FIG. 12.**
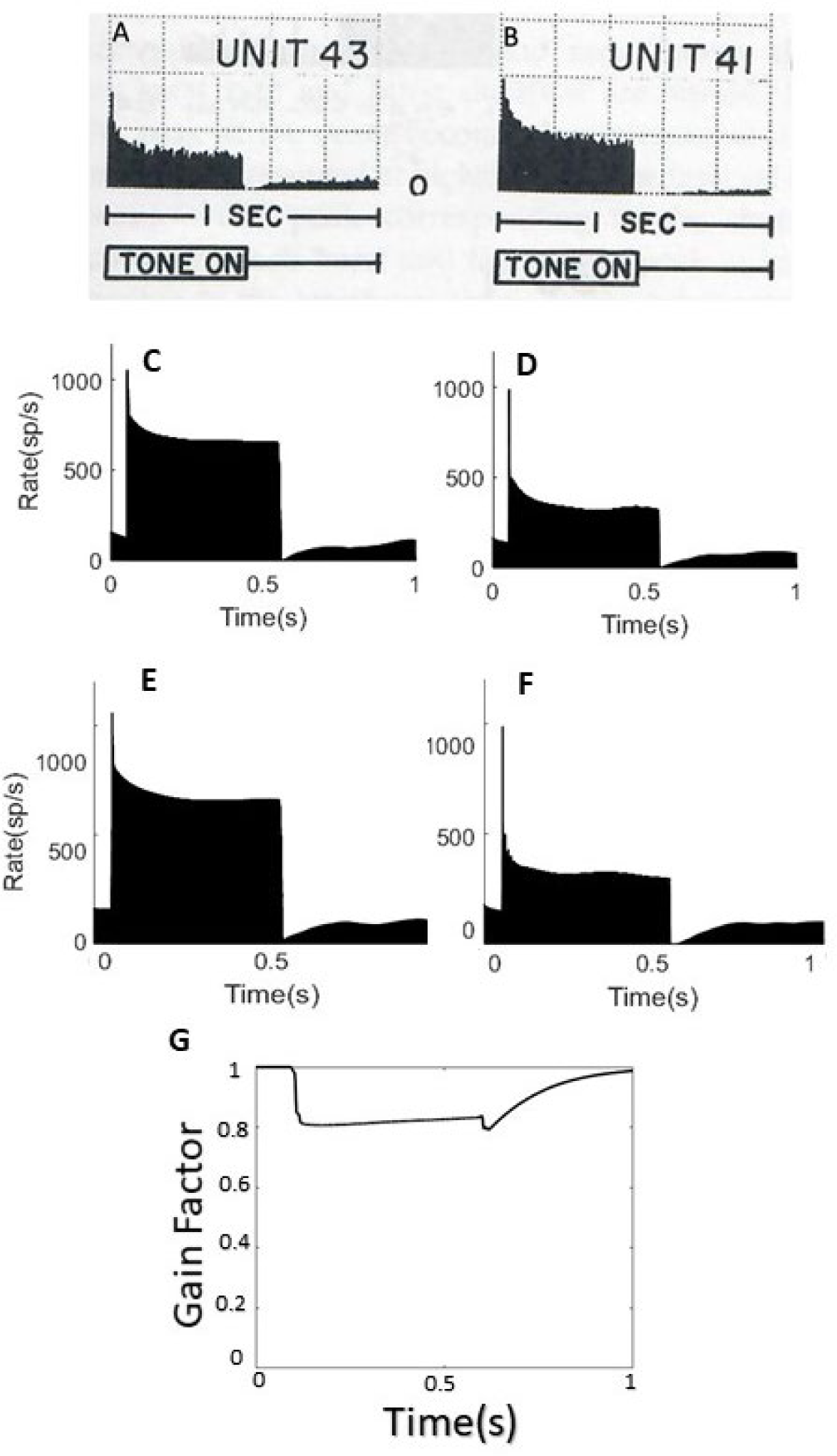
AN and model responses to tones at CF: A) Unit 43: CF = 1.82 kHz, 55 dB SPL, SR = 60.6 s/sec; B) Unit 41: CF = 10.34 kHz, 30 dB SPL, SR = 51.7 s/sec; Panels A, B adapted from Kiang, Nelson Yuan-Sheng. with the assistance of Takeshi Watanabe, Eleanor C. Thomas, and Louise F. Clark., Discharge Patterns of Single Fibers in the Cat’s Auditory Nerve, Fig 6.1 (p. 69), © 1966 Massachusetts Institute of Technology, by permission of The MIT Press. C, D) Responses to the same tones for the model without efferents, and E, F) with MOC efferents. G) The gain factor variation over time for the model with efferents (gain factor for plot E). For all model simulations (C to F) the AN model type was HSR with CFs matching the AN fibers in the physiological study. The stimulus sound level was matched to stimuli for the physiological data.

#### Model response to an increment in tone level

One of the paradigms tested in Zilany et al. (2009) was the model response to an increment in tone level. In the physiological data (Westerman & Smith, 1987) the AN fiber response to a tone at the fiber’s CF (5.99 kHz, HSR) was recorded in anesthetized cat. Over the first 300 ms of the stimulus, the tone level was 5, 10, 15, or 20 dB above threshold, and for the second 300 ms, the tone level increased to 43 dB above threshold. In the physiological responses, the transient response to the pedestal increased with pedestal level, but the transient response to the increment decreased with increasing pedestal level. The same trend was observed in the response of the AN model with and without MOC efferents. The increase in rate over time that followed the first transient response was observed for the model with MOC efferents (Fig. 13C, for initial tone levels of +5 and +10 dB), but not for physiological responses in the anesthetized gerbil (Fig. 13A). This discrepancy between the MOC model predictions and the physiological data may be due to the effects of anesthesia on the empirical AN recordings. The increase in rate during the low-level tones (Fig. 13-c, +5 and +10 dB) reflects the recovery of cochlear gain after an initial decrease in response to the onset of the stimulus. Responses in unanesthetized animals are not available for these stimuli, therefore, it is not possible to confirm this interpretation at this time.

**FIG. 13.**
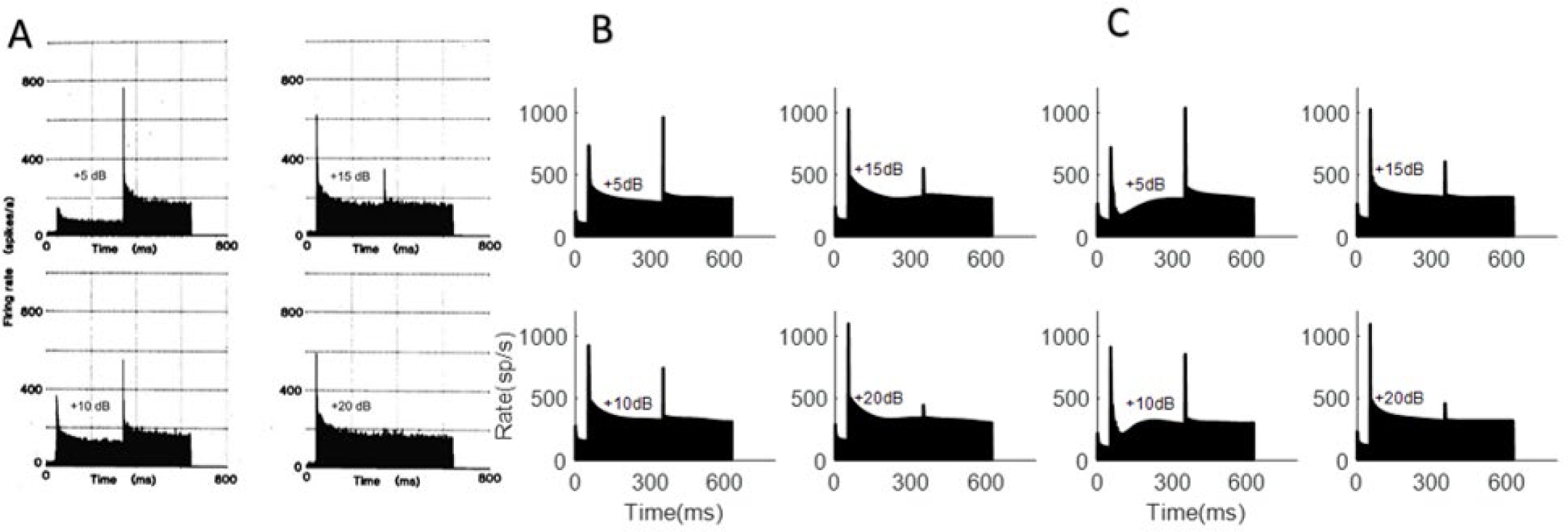
A) Physiological data from gerbil (adapted from Westerman and Smith (1987) with permission of Acoustical Society of America) in response to an increase in level during a tone. The AN model response to the amplitude increment response paradigm B) without efferent system C) with MOC efferent system.

#### Model response to AM stimuli

Here the model response was compared to the response of an HSR AN fiber with CF of 20.2 kHz to an AM tone with a carrier frequency at CF (Joris & Yin, 1992). The effect of modulation depth on the synchrony to modulation frequency (Fm) was studied for modulation depths ranging from 0 to 1 (Fig. 14A). The modulation frequency was 100 Hz and level was 49 dB SPL. A comparison of synchronization to AM frequency as a function of modulation depth showed that the model with MOC efferents had increased synchrony with modulation depth. However, the synchrony of the model response was weaker than that observed physiologically because of the reduction in cochlear gain as a result of MOC efferent activity. Complications for comparison exist due to the difference between dynamic ranges of the model and this physiological example, as noted by Zilany et al., (2009). Furthermore, model responses to AM tones, which consist of both carrier and sideband frequencies, may be shaped by the effective bandwidths of MOC projections, discussed below. AM responses of the model with efferents, however, cannot be directly compared to physiological responses, which are not available for AN fibers in unanesthetized animals. However, we do have access to IC responses to AM noise in awake rabbit. Figure 14B compares the synchrony to AM noise for the IC stage of the model with MOC efferents to actual responses from a single-unit recording in the IC of awake rabbit (Carney et al., 2014). This result confirms that, for the model with efferents, the increase in synchrony in the AN model with increasing modulation depth results in appropriate synchrony at the level of the model IC.

**FIG. 14.**
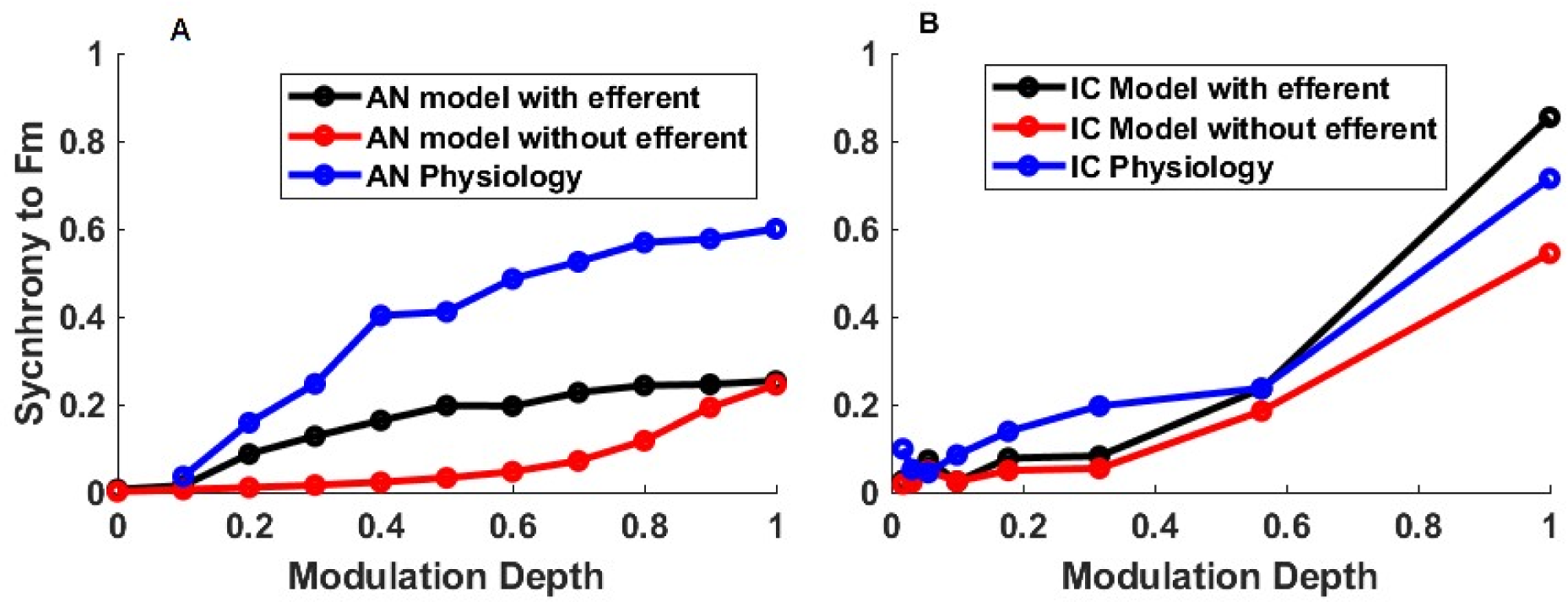
(color online) A) Synchrony to the envelope of an AM tone at CF for an AN fiber with CF = 20 kHz (AN Physiology data from Joris & Yin, 1992) and for an HSR model fiber with matching CF with and without efferent activation. Modulation frequency of 100 Hz; level = 49 dB SPL. B) IC model with and without activation of the MOC efferent system and IC physiology synchronization-modulation depth plots for responses to AM noise stimulus at CF 5500 Hz, BMF 100 Hz, modulation frequency 64 Hz and stimulus level 36 dB SPL.

## IV. DISCUSSION

In this work, we modified an existing model of the peripheral auditory system to simulate both the ascending pathway and MOC efferent feedback with both CN and IC inputs. The physiological data and simulation results are consistent with the hypothesis that the MOC efferent system influences neural fluctuations over time (Carney, 2018), as observed here by the increasing responses of BE IC cells over the time course of an AM noise. The simulation results support the hypothesis that a sub-cortical model with MOC efferent gain control better simulates IC responses to AM noise compared to a model without efferents.

One of the components of the proposed model is a LPF that was implemented to simulate the dynamics of the MOC efferent system. The time constant (or cut-off frequency) of this filter was adjusted based on the physiological dataset. The dynamics of the MOC efferent system based on the fit parameter agree with the reported dynamics in the literature for the MOC efferent fast effect. Backus and Guinan (2006) used stimulus-frequency otoacoustic emission (SFOAE) measurements to estimate the human MOC-reflex time constant. They reported an overall onset time constant of 277±62 ms and decay time constant of 159±54 ms. Warren and Liberman (1989) estimated the MOC response to have a time constant of 250 ms in barbiturate-anesthetized cats. Roverud and Strickland (2010) used an optimized psychophysical technique to estimate the time course of cochlear-gain reduction. They estimated an average exponential decay of 100 ms, which is faster than the time constant estimated by Backus and Guinan. Yasin et al. (2020) used a computational model of human hearing that included MOC efferent feedback and evaluated the model’s speech recognition for different MOC efferent time constants. Their results suggest the improvement in recognition of speech in AM noise is highest for a MOC efferent time constant of 200 ms. In general, all of the time constants suggested in the literature for the MOC efferent system are in the same range as our results based on rabbit midbrain physiology. However, the MOC efferent system time constant could differ between rabbits and humans. Using the proposed MOC efferent model to simulate human psychoacoustic experiments could better inform our understanding of the time constant for human listeners.

The dynamics of the efferent system support its potential role in improving speech perception (Yasin et al., 2018). Specifically, the latency and time constant of the MOC efferent system (Backus & Guinan, 2006; Roverud & Strickland, 2010; Warren III & Liberman, 1989) are comparable to typical phonemic durations (Jacewicz et al., 2007; Umeda, 1977). Therefore, the MOC efferent system would presumably affect auditory neural responses, both within the time course of phonemes and also in running speech, for which the perception of a phoneme would be influenced by the previous phoneme (co-articulation) (Chambers et al., 2017; Recasens, 1984, 2018).

Another important component of the proposed model MOC efferent system was the input-output function that maps the MOC input to a gain factor that scales the cochlear-gain coefficient between 0 and 1. The second-order rational function used in this model was chosen based on its asymptotic behavior at low and high input levels. The direct input-output function of MOC neurons is not known, but several related functions have been reported. The shape of the proposed input-output function was similar to the MOC efferent attenuation as a function of input sound level shown in Russell and Murugasu (1997). Also, the asymptotic trend of this function for higher values is similar to the asymptotic responses shown in Liberman (1988) for MOC efferent activity as a function of input sound level. Finally, the proposed function is similar to the relationship between stimulus level and OHC power gain, which would be modulated by MOC signals, in an electrical-mechanical model for OHCs (Rabbitt & Bidone, 2023; Rabbitt, 2023).

### A. Comparing the Proposed model with previous AN models that included the efferent system

Several previous AN models include efferent signals driven by the periphery (Clark et al., 2012; Giguere & Woodland, 1994; Kwan et al., 2019; Smalt et al., 2014; Yasin et al., 2020). In some of these previous implementations, cochlear gain was set to remain constant over time, omitting the dynamics of MOC efferents (Brown et al., 2010; Ferry & Meddis, 2007; Giguere & Woodland, 1994; Jennings et al., 2011). Smalt et al. (2014) proposed a model based on the Zilany et al. (2014) AN model, but with an efferent system modeled by a time-varying gain-control feedback system. However, in this model, a local gain-control system was implemented based on the output of model OHCs, which is inconsistent with the overall structure of the MOC efferent system, which depends on outputs from the OHCs, IHC, AN, CN, IC, and other structures (Smalt et al., 2014; Schofield, 2011). None of these models simulated the descending input to the MOC efferent system from the midbrain. The focus of the current study was to include both peripheral and midbrain inputs to a dynamic MOC efferent system in a model for AN and BE IC responses.

### B. Limitations and Future directions

Other physiological and psychoacoustic studies can be used to further fine tune the model’s parameters and structure. For example, forward-masking studies with long-duration maskers would be expected to activate the MOC efferent system (Krull and Strickland, 2008; Brennan et al., 2023). At the level of the IC, further animal studies of the temporal properties of responses to complex sounds would contribute to this modeling effort, such as investigating the changes in synchrony of IC responses to the envelope of AM stimuli over time. Another direction for future work is to study the influence of efferent activity on the responses of other MTF types in the IC, such as BS or hybrid MTFs. Here we focused on BE MTFs, as the increase in their rate over the time course of AM stimuli cannot be confused with rate adaptation. Phase of AM coding could be an important aspect for studying speech. AM phase was not explored in this study due to the relatively short stimulus duration used and the limited number of repetitions of the stimuli in the available dataset. Physiological experiments designed to study the effect of the efferent system on the phase of AM coding is an interesting future direction.

The peripheral model studied here is a monaural model, with “one-sided” brainstem and midbrain stages. A binaural model would be a powerful tool for studying the crossed and uncrossed effects of the efferents. The proposed model in this study consists of a single CF channel, and MOC controls for each frequency influence the gain of the related CF channel. However, studies have shown that some projections from MOC neurons to OHCs have a broad distribution across CF channels (Brown, 2014; Lee & Lewis, 2023). To move towards a more comprehensive model, a future direction of research is to explore a cross-frequency projection of the MOC control system that affects a range of CFs. A broader distribution of MOC projections could benefit simulations of the neural response to complex stimuli, such AM noise and also speech signals. Another direction for future work on this model is inclusion of a more advanced IC model, instead of the simple bandpass filter used for modulation filtering in BE IC neurons in the current model.

In general, recording from AN fibers requires an anesthetized animal preparation, but anesthesia suppresses MOC efferent activity (Guitton et al., 2004). As a result, there are no examples in the AN physiological literature for comparison to the AN model response with active MOC efferents. However, recording from more accessible levels of the auditory pathway, such as the IC in awake animals, could provide further information regarding the MOC efferent system. Recording from the CN in awake animals or recording from AN fibers in decerebrate animals could also potentially be beneficial for studying the MOC efferent system (Kim et al., 1990a, 1990b; Kawase et al., 1993).

Other phenomena could explain the increase in IC response rates over time in response to AM noise, either in addition to or instead of efferent activity. For instance, adaptation of inhibition (Viemeister & Bacon, 1982; Oberle et al., 2023) could be an alternative explanation for the increase in rate we observed in IC responses to AM noise. Adaptation of inhibition has been suggested as an explanation for auditory enhancement, in which a target signal embedded in a masker is preceded by the same masker but without the target frequency. Adaptation of inhibition could explain an improvement in the threshold of detecting the tone (Byrne et al., 2011). However, neural mechanisms underlying adaptation of inhibition are not clear. Enhancement effects and its properties, such as time constant and level-dependence, may be further examined using the MOC efferent model in the future to see if such effects are consistent with MOC feedback. Preliminary results suggest that the subcortical model with efferent feedback can also explain several aspects of auditory enhancement (Farhadi et al., 2023a).

In addition to subcortical projections, there are significant descending projections from the auditory cortex and thalamus to the IC (Schofield, 2011). The descending projection to the central nucleus of the IC (ICc) could potentially contribute to the observed trend in the physiology dataset. However, studies indicate that in animal models (except for rats), the projections from the auditory cortex and medial geniculate nucleus (MG) in the thalamus terminate in the external cortex of the IC (ICx) and dorsal cortex of the IC (ICd), and the projection to ICc is described as much less dense than ICd or ICx (Schofield, 2011). These projections were not included in the proposed subcortical model, but they may influence responses of IC cells.

## V. ACKNOWLEDGMENTS

This work was supported by NIH-R01-DC010813 (AF, LHC), NIH-R01-DC008327 (EAS), and K23 DC014752 (SGJ).

## VI. AUTHOR DECLARATIONS

The authors have no conflicts of interest to disclose.

## VII. DATA AVAILABILITY

Model code will be made freely available as downloadable files (Farhadi et al., 2023) from the Open Science Framework, DOI 10.17605/OSF.IO/UZYW4.

## REFERENCES

Aedo, C., Tapia, E., Pavez, E., Elgueda, D., Delano, P. H., & Robles, L. (2015). “Stronger efferent suppression of cochlear neural potentials by contralateral acoustic stimulation in awake than in anesthetized chinchilla,” Frontiers in Systems Neuroscience 9, 21.

Backus, B. C., & Guinan Jr, J. J. (2006). “Time-course of the human medial olivocochlear reflex,” J. Acoust. Soc. Am. 119, 2889–2904.

Bharadwaj, H. M., Verhulst, S., Shaheen, L., Liberman, M. C., & Shinn-Cunningham, B. G. (2014). “Cochlear neuropathy and the coding of supra-threshold sound,” Frontiers in Systems Neurosci. 8, 26.

Blackburn, C. C., & Sachs, M. B. (1990). “The representations of the steady-state vowel sound/e/in the discharge patterns of cat anteroventral cochlear nucleus neurons,” J. Neurophys. 63, 1191–1212.

Brennan, M. A., Svec, A., Farhadi, A., Maxwell, B. N., & Carney, L. H. (2023). “Inherent envelope fluctuations in forward masking: Effects of age and hearing loss,” J. Acoust. Soc. Am. 153, 1994–2005.

Brown, M. C. (2014). “Single-unit labeling of medial olivocochlear neurons: the cochlear frequency map for efferent axons,” J. Neurophys. 111, 2177–2186.

Brown, G. J., Ferry, R. T., & Meddis, R. (2010). “A computer model of auditory efferent suppression: Implications for the recognition of speech in noise,” J. Acoust. Soc. Am. 127, 943–954.

Byrne, A. J., Stellmack, M. A., & Viemeister, N. F. (2011). “The enhancement effect: evidence for adaptation of inhibition using a binaural centering task,” J. Acoust. Soc. Am. 129, 2088–2094.

Carney, L. H. (2018). “Supra-threshold hearing and fluctuation profiles: implications for sensorineural and hidden hearing loss,” JARO 19, 331–352.

Carney, L. H., Li, T., & McDonough, J. M. (2015). “Speech coding in the brain: representation of vowel formants by midbrain neurons tuned to sound fluctuations,” Eneuro 2, 4.

Carney, L. H., Zilany, M. S. A., Huang, N. J., Abrams, K. S., & Idrobo, F. (2014). “Suboptimal use of neural information in a mammalian auditory system,” J. Neurosci. 34, 1306–1313.

Chambers, A. R., Hancock, K. E., Maison, S. F., Liberman, M. C., & Polley, D. B. (2012). “Sound-evoked olivocochlear activation in unanesthetized mice,” JARO 13, 209–217.

Chambers, C., Akram, S., Adam, V., Pelofi, C., Sahani, M., Shamma, S., & Pressnitzer, D. (2017). “Prior context in audition informs binding and shapes simple features,” Nat. Comm. 8, 1–11.

Clark, N. R., Brown, G. J., Jürgens, T., & Meddis, R. (2012). “A frequency-selective feedback model of auditory efferent suppression and its implications for the recognition of speech in noise,” J. Acoust. Soc. Am. 132, 1535–1541.

Delgutte, B., & Kiang, N. Y. (1984). “Speech coding in the auditory nerve: I. Vowel-like sounds,” J. Acoust. Soc. Am. 75, 866–878.

Delano, P. H., Elgueda, D., Hamame, C. M., & Robles, L. (2007). “Selective attention to visual stimuli reduces cochlear sensitivity in chinchillas,” J. Neurosci. 27, 4146–4153.

Deng, L., Geisler, C. D., & Greenberg, S. (1987). “Responses of auditory-nerve fibers to multiple-tone complexes,” J. Acoust. Soc. Am. 82, 1989–2000.

Farhadi, A., Agarwalla, S. & Carney, L. H. (2023a). “A Subcortical Auditory Model With Efferent Gain Control Explains Perceptual Enhancement,” Abstract, Assoc. Res. Otolaryngol. 46:449.

Farhadi, A., Jennings, S. G., Strickland, E. A., & Carney, L. H. (2023b). Downloadable MATLAB/C Code for “Subcortical Auditory Model including Efferent Dynamic Gain Control with Inputs from Cochlear Nucleus and Inferior Colliculus” from OSF: https://osf.io/uzyw4/, DOI 10.17605/OSF.IO/UZYW4, Software.

Farhadi, A., Jennings, S. G., Strickland, E. A., & Carney, L. H. (2021). “A Closed-Loop Gain-Control feedback model for the medial efferent system of the descending auditory pathway,” In ICASSP 2021-2021 IEEE Intl. Conf. Acoust. Speech Signal Proc. (ICASSP) (pp. 291–295).

Fekete, D. M., Rouiller, E. M., Liberman, M. C., & Ryugo, D. K. (1984). “The central projections of intracellularly labeled auditory nerve fibers in cats,” J. Comp. Neurol. 229, 432–450.

Ferry, R. T., & Meddis, R. (2007). “A computer model of medial efferent suppression in the mammalian auditory system,” J. Acoust. Soc. Am. 122, 3519–3526.

Giguere, C., & Woodland, P. C. (1994). “A computational model of the auditory periphery for speech and hearing research. II. Descending paths,” J. Acoust. Soc. Am. 95, 343–349.

Guinan Jr, J. J. (2006). “Olivocochlear efferents: anatomy, physiology, function, and the measurement of efferent effects in humans,” Ear Hear. 27, 589–607.

Guinan Jr, J. J. (2018). “Olivocochlear efferents: Their action, effects, measurement and uses, and the impact of the new conception of cochlear mechanical responses,” Hear. Res. 362, 38–47.

Guitton, M. J., Avan, P., Puel, J.-L., & Bonfils, P. (2004). “Medial olivocochlear efferent activity in awake guinea pigs,” Neuroreport 15, 1379–1382.

Gummer, M., Yates, G. K., & Johnstone, B. M. (1988a). “Modulation transfer function of efferent neurones in the guinea pig cochlea,” Hear. Res. 36, 41–51.

Gummer, M., Yates, G. K., & Johnstone, B. M. (1988b). “Modulation transfer function of efferent neurones in the guinea pig cochlea,” Hear. Res. 36, 41–51.

Huffman, R. F., & Henson Jr, O. W. (1990). “The descending auditory pathway and acousticomotor systems: connections with the inferior colliculus,” Brain Res. Rev. 15, 295–323.

Jacewicz, E., Fox, R. A., & Salmons, J. (2007). “Vowel duration in three American English dialects,” American Speech 82, 367–385.

Jamos, A. M., Kaf, W. A., Chertoff, M. E., & Ferraro, J. A. (2020). “Human medial olivocochlear reflex: contralateral activation effect on low and high frequency cochlear response,” Hear. Res. 389, 107925.

Jamos, A. M., Hosier, B., Davis, S., & Franklin, T. C. (2021). “The role of the medial olivocochlear reflex in acceptable noise level in adults,” J. Am. Acad. Audiol. 32, 137–143.

Jennings, S. G., & Aviles, E. S. (2023). “Middle ear muscle and medial olivocochlear activity inferred from individual human ears via cochlear potentials.” J. Acoust. Soc. Am. 153, 1723–1732.

Jennings, S. G. (2021). “The role of the medial olivocochlear reflex in psychophysical masking and intensity resolution in humans: a review,” J. Neurophys. 125, 2279–2308.

Jennings, S. G., Heinz, M. G., & Strickland, E. A. (2011). “Evaluating adaptation and olivocochlear efferent feedback as potential explanations of psychophysical overshoot,” JARO 12, 345–360.

Joris, P. X., Schreiner, C. E., & Rees, A. (2004). “Neural processing of amplitude-modulated sounds,” Phys. Rev. 84, 541–577.

Joris, P. X., & Yin, T. C. T. (1992). “Responses to amplitude-modulated tones in the auditory nerve of the cat,” J. Acoust. Soc. Am. 91, 215–232.

Kawase, T., Delgutte, B., & Liberman, M. C. (1993). “Antimasking effects of the olivocochlear reflex. II. Enhancement of auditory-nerve response to masked tones,” J. Neurophys. 70, 2533–2549.

Kiang, N. Y. S., Watanabe, T., Thomas, E. C., & Clark, L. F. (1965). Discharge patterns of single fibers in the cat’s auditory nerve. Cambridge, MA: MIT Press.

Kim, D. O., Carney, L., & Kuwada, S. (2020). “Amplitude modulation transfer functions reveal opposing populations within both the inferior colliculus and medial geniculate body,” J. Neurophys. 124, 1198–1215.

Kim, D. O., Zahorik, P., Carney, L. H., Bishop, B. B., & Kuwada, S. (2015). “Auditory distance coding in rabbit midbrain neurons and human perception: monaural amplitude modulation depth as a cue,” J. Neurosci. 35, 5360–5372.

Kim, D. O., Sirianni, J. G., & Chang, S. O. (1990). “Responses of DCN-PVCN neurons and auditory nerve fibers in unanesthetized decerebrate cats to AM and pure tones: analysis with autocorrelation/power-spectrum,” Hear. Res. 45, 95–113.

Kim, D. O., Chang, S. O., & Sirianni, J. G. (1990). “A population study of auditory-nerve fibers in unanesthetized decerebrate cats: Response to pure tones,” J. Acoust. Soc. Am. 87, 1648–1655.

Krishna, B. S., & Semple, M. N. (2000). “Auditory temporal processing: responses to sinusoidally amplitude-modulated tones in the inferior colliculus,” J. Neurophys. 84, 255–273.

Krull, V., & Strickland, E. A. (2008). “The effect of a precursor on growth of forward masking,” J. Acoust. Soc. Am. 123, 4352–4357.

Kwan, T. J. M., Zilany, M. S. A., Davies-Venn, E., & Abdul Wahab, A. K. (2019). “Modeling the effects of medial olivocochlear efferent stimulation at the level of the inferior colliculus,” Exp. Brain Res. 237, 1479–1491.

Lehky, S. R. (2010). “Decoding poisson spike trains by gaussian filtering,” Neural Comp. 22, 1245–1271.

Lauer, A. M., Jimenez, S. V., & Delano, P. H. (2022). “Olivocochlear efferent effects on perception and behavior,” Hear. Res. 419, 108207.

Lee, D., & Lewis, J. D. (2023). “Inter-Subject Variability in the Dependence of Medial-Olivocochlear Reflex Strength on Noise Bandwidth,” Ear Hear. 44, 544–557.

Liberman, M. C. (1988). “Response properties of cochlear efferent neurons: monaural vs. binaural stimulation and the effects of noise,” J. Neurophys. 60, 1779–1798.

Liberman, M. C. (1978). “Auditory-nerve response from cats raised in a low-noise chamber,” J. Acoust. Soc. Am. 63, 442–455.

Liberman, M. C., & Guinan Jr, J. J. (1998). “Feedback control of the auditory periphery: antimasking effects of middle ear muscles vs. olivocochlear efferents,” J. Comm. Dis. 31, 471–482.

Lichtenhan, J. T., Wilson, U. S., Hancock, K. E., & Guinan Jr, J. J. (2016). “Medial olivocochlear efferent reflex inhibition of human cochlear nerve responses,” Hear Res. 333, 216–224.

Lopez-Poveda, E. A. (2018). “Olivocochlear efferents in animals and humans: from anatomy to clinical relevance,” Frontiers Neurol. 9, 197.

Mao, J., Vosoughi, A., & Carney, L. H. (2013). “Predictions of diotic tone-in-noise detection based on a nonlinear optimal combination of energy, envelope, and fine-structure cues,” J. Acoust. Soc. Am. 134, 396–406.

May, B. J., & Sachs, M. B. (1992). “Dynamic range of neural rate responses in the ventral cochlear nucleus of awake cats,” J. Neurophys. 68, 1589–1602.

Nelson, P. C., & Carney, L. H. (2004). “A phenomenological model of peripheral and central neural responses to amplitude-modulated tones,” J. Acoust. Soc. Am. 116, 2173–2186.

Oberle, H. M., Ford, A. N., Czarny, J. E., Rogalla, M. M., & Apostolides, P. F. (2023, in press). “Recurrent circuits amplify corticofugal signals and drive feedforward inhibition in the inferior colliculus,” J. Neurosci.

Oertel, D., Wright, S., Cao, X.-J., Ferragamo, M., & Bal, R. (2011). “The multiple functions of T stellate/multipolar/chopper cells in the ventral cochlear nucleus,” Hear. Res. 276, 61–69.

Olsen, W. O. (1998). “Average speech levels and spectra in various speaking/listening conditions,” Am. J. Audiol. 7, 21–25.

Rabbitt, R. (2023) “The MOC efferent system acts as a control parameter for outer hair cell dynamic stability and power output,” Abstract, Assoc. Res. Otolaryngol. 46, 544.

Rabbitt, R., and Bidone, T.C. (2023). “A parametric blueprint for optimum cochlear outer hair cell design,” J. Roy. Soc. Interface 20, 20220762.

Rahman, M., Willmore, B. D. B., King, A. J., & Harper, N. S. (2019). “A dynamic network model of temporal receptive fields in primary auditory cortex,” PLoS Comp. Biol. 15, e1006618.

Recasens, D. (1984). “Vowel-to-vowel coarticulation in Catalan VCV sequences,” J. Acoust. Soc. Am. 76, 1624–1635.

Recasens, D. (2018). “Coarticulation,” In Oxford Research Encyclopedia of Linguistics. 10.1093/acrefore/9780199384655.013.416.

Rhode, W. S., & Greenberg, S. (1994). “Encoding of amplitude modulation in the cochlear nucleus of the cat,” J. Neurophys. 71, 1797–1825.

Rhode, W. S., & Kettner, R. E. (1987). “Physiological study of neurons in the dorsal and posteroventral cochlear nucleus of the unanesthetized cat,” J. Neurophys. 57, 414–442.

Romero, G. E., & Trussell, L. O. (2021). “Distinct forms of synaptic plasticity during ascending vs descending control of medial olivocochlear efferent neurons,” Elife 10, e66396.

Roverud, E., & Strickland, E. A. (2010). “The time course of cochlear gain reduction measured using a more efficient psychophysical technique,” J. Acoust. Soc. Am. 128, 1203–1214.

Russell, I. J., & Murugasu, E. (1997). “Medial efferent inhibition suppresses basilar membrane responses to near characteristic frequency tones of moderate to high intensities,” J. Acoust. Soc. Am. 102, 1734–1738.

Ryugo, D. K. (2008). “Projections of low spontaneous rate, high threshold auditory nerve fibers to the small cell cap of the cochlear nucleus in cats,” Neurosci. 154, 114–126.

Schofield, B. R. (2011). “Central descending auditory pathways,” In Auditory and vestibular efferents (pp. 261–290). Springer.

Shera, C. A., Guinan Jr, J. J., & Oxenham, A. J. (2002). “Revised estimates of human cochlear tuning from otoacoustic and behavioral measurements,” PNAS 99, 3318–3323.

Smalt, C. J., Heinz, M. G., & Strickland, E. A. (2014). “Modeling the time-varying and level-dependent effects of the medial olivocochlear reflex in auditory nerve responses,” JARO 15, 159–173.

Smith, S. B., Lichtenhan, J. T., & Cone, B. K. (2017). “Contralateral inhibition of click-and chirp-evoked human compound action potentials,” Frontiers Neurosci. 11, 189.

Sun, X.-M. (2008). “Contralateral suppression of distortion product otoacoustic emissions and the middle-ear muscle reflex in human ears,” Hear. Res. 237, 66–75.

Tan, Q., & Carney, L. H. (2003). “A phenomenological model for the responses of auditory-nerve fibers. II. Nonlinear tuning with a frequency glide,” J. Acoust. Soc. Am. 114, 2007–2020.

Umeda, N. (1977). “Consonant duration in american english,” J. Acoust. Soc. Am. 61, 846–858.

Viemeister, N. F., & Bacon, S. P. (1982). “Forward masking by enhanced components in harmonic complexes,” J. Acoust. Soc. Am. 71, 1502–1507.

Warr, W. B. (1992). “Organization of olivocochlear efferent systems in mammals,” In The Mammalian Auditory Pathway: Neuroanatomy, pp. 410–448. Springer.

Warren III, E. H., & Liberman, M. C. (1989). “Effects of contralateral sound on auditory-nerve responses. I. Contributions of cochlear efferents,” Hear. Res. 37, 89–104.

Westerman, L. A., & Smith, R. L. (1984). “Rapid and short-term adaptation in auditory nerve responses,” Hear. Res. 15, 249–260.

Westerman, L. A., & Smith, R. L. (1987). “Conservation of adapting components in auditory-nerve responses,” J. Acoust. Soc. Am. 81, 680–691.

Yasin, I., Drga, V., Liu, F., Demosthenous, A., & Meddis, R. (2020). “Optimizing Speech Recognition Using a Computational Model of Human Hearing: Effect of Noise Type and Efferent Time Constants,” IEEE Access 8, 56711–56719.

Yasin, I., Liu, F., Drga, V., Demosthenous, A., & Meddis, R. (2018). “Effect of auditory efferent time-constant duration on speech recognition in noise,” J. Acoust. Soc. Am. 143, EL112–EL115

Ye, Y., Machado, D. G., & Kim, D. O. (2000). “Projection of the marginal shell of the anteroventral cochlear nucleus to olivocochlear neurons in the cat,” J. Comp. Neurol. 420, 127–138.

Young, E. D., & Sachs, M. B. (1979). “Representation of steady-state vowels in the temporal aspects of the discharge patterns of populations of auditory-nerve fibers,” J. Acoust. Soc. Am. 66, 1381–1403.

Zhang, X., Heinz, M. G., Bruce, I. C., & Carney, L. H. (2001). “A phenomenological model for the responses of auditory-nerve fibers: I. Nonlinear tuning with compression and suppression.” J. Acoust. Soc. Am. 109, 648–670.

Zilany, M. S. A., & Bruce, I. C. (2006). “Modeling auditory-nerve responses for high sound pressure levels in the normal and impaired auditory periphery,” J. Acoust. Soc. Am. 120, 1446–1466.

Zilany, M. S. A., Bruce, I. C., & Carney, L. H. (2014). “Updated parameters and expanded simulation options for a model of the auditory periphery,” J. Acoust. Soc. Am. 135, 283–286.

Zilany, M. S. A., Bruce, I. C., Nelson, P. C., & Carney, L. H. (2009). “A phenomenological model of the synapse between the inner hair cell and auditory nerve: long-term adaptation with power-law dynamics,” J. Acoust. Soc. Am. 126, 2390–2412.

